# Discovery of intratumoral oncolytic bacteria toward targeted anticancer theranostics

**DOI:** 10.1101/2022.10.25.513676

**Authors:** Yamato Goto, Seigo Iwata, Eijiro Miyako

## Abstract

Unveiling the different biomedical functions of tumor-resident microbiota has remained challenging for the development of advanced anticancer medicines. Here we show that isolated intratumoral bacteria with its association with natural purple photosynthetic bacteria have a high innate biocompatibility and drastic immunogenic anticancer efficacies. They preferentially grow and proliferate within targeted tumor milieu, which effectively causes immune cells to infiltrate the tumor and provoke strong anticancer responses in various syngeneic mouse models including those of colorectal cancer, sarcoma, metastatic lung cancer, and extensive drug-resistant breast cancer. Furthermore, these functional bacteria-treated mice, that exhibit excellent anticancerous responses of tumors, have significantly prolonged survival rates with effective immunological memory. Notably, light-harvesting nanocomplexes of microbial consortium of intratumoral bacteria and purple photosynthetic bacteria is capable of tumor diagnosis using bio-optical-window near-infrared light, making them useful theranostic agents for highly targeted immunological elimination of the tumor and for precisely marking tumor location.

## Introduction

Advanced anticancer medicines, that are highly effective against targeted tumors, are multifunctional, simplistic, safe, and inexpensive. Such treatments are essential for patients who experience medicinal side effects and financial issues in typical cancer remedies^1^. Bacterial cancer therapies, employed in clinical trials, are substantially attractive for targeted cancer elimination because bacteria have promising properties that make them viable candidates for the treatment^2–6^. Although bacteria are fashionable as a therapeutic agent, conventional cancer therapy typically requires genetic engineering techniques^7, 8^, synthetic bioengineering^9,10^, and nanotechnology^1^^1–13^ for bacterial attenuation and improved drug efficacy because bacteria which were originally pathogenic and have low medicinal properties are used. Moreover, various natural bacteria have been unaccountably applied for potential cancer treatment by many researchers for more than a century^2–6^. Nevertheless, we believe that it is important to further explore biocompatible natural bacteria which have innate stronger efficacies and higher tumor specificity, without necessitating genetic manipulations and nano engineering^14^.

Tumor-resident microbiota may be another approach for exploration of novel functional anticancer natural bacterial therapy. Recent evidence suggests that intratumoral bacteria can influence the efficacy of cancer therapies^15, 16^. However, most studies have mainly focused on a causal role of intratumoral bacteria in the development of cancer or on their presence reflects anticancer efficacies of chemotherapy against tumors^17–19^. Also, intratumoral bacteria themselves have not been used as an anticancer therapeutic agent directly, although there is an unrevealed possibility in their use for cancer treatment.

Here, we aimed to explore highly targeted cancer immunotherapeutic bacteria from the tumor-resident microbiota and its association with natural photosynthetic bacteria. Further, we assessed their functional anticancer efficacies such as tumor targeting ability and biocompatibility in various syngeneic mouse models regardless of the presence or absence of immunogenicity in tumor, and analysed the survival rates of mice with unique immunological memory. In a targeted tumor theranostic approach, unique optical properties of intratumoral bacteria and photosynthetic bacteria was also explored using tissue-penetrating near-infrared (NIR) light. This research will shed light on the ever evolving biology of tumors and provide a novel approach to the treatment of cancer in addition to developments of advanced optical precision medical devices.

## Results

### Isolation of highly effective anticancer bacteria from tumors

Previously, we found that non-pathogenic natural purple photosynthetic bacteria *Rhodopseudomonas palustris* (RP) and *Blastochloris viridis* displayed multifunctionality and biocompatibility as cancer theranostic agents in the treatment of highly active cancers, using bio-optical-window I and II NIR light^14^. A major challenge was the exploration of functional photosynthetic bacteria with superior anticancer therapeutic efficacy which were accidentally discovered during the extractions of purple photosynthetic bacteria RP from its infected tumor biopsy for further characterisations. **Figure 1** depicts a schematic illustration of isolation of extremely effective anticancer bacteria from a solid tumor. RP can colonise and grow in a tumor with high specificity because it prefers hypoxic environment^13^. We have previously reported characteristic red colonies of RP which were formed on an ATCC 543 agar plate after streaking the solution extracted from the RP-infected tumor. Although conventional RP suspension naturally reveals red colour, the extracted solution from a solid tumor unexpectedly became grey in colour over time because of the contamination with tumor-resident microbiota^17, 18^. *Proteus mirabilis* (PM) is anaerobic, gram-negative, and a member of the intratumoral bacterial family *Enterobacteriaceae*^20, 21^. Many white circular colonies of PM were formed on ATCC 543 agar plate with sodium deoxycholate (SD) that was used to culture single colonies as SD prohibits bacterial swarming (migration across surfaces of solid media) of PM^22^. Isolated intratumoral PM (i-PM) was confirmed to be 99.74% pure using gene identification (**Supplementary Table S1**).

**Figure 1.**
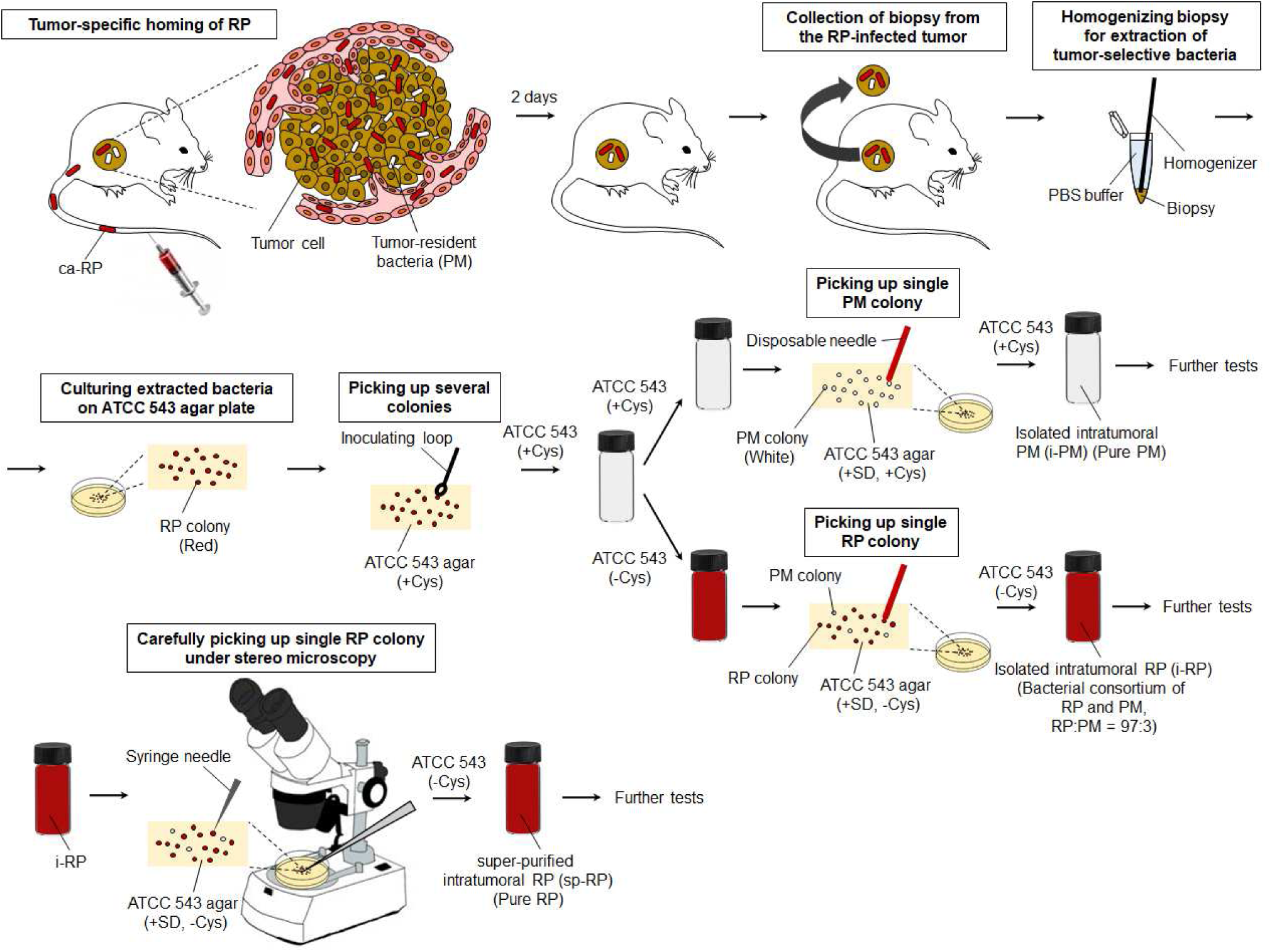
Schematic illustration of isolation of extremely effective anticancer bacteria from solid tumors.

Meanwhile, the solution of bacteria in the ATCC 543 medium without L-cysteine (Cys) turned red after subculturing, and provided red and white colonies on an ATCC 543 agar plate with SD and without Cys. Although SD restraints bacterial swarming of PM, indistinct and translucent colonies of PM intermingle with RP colonies thus making it difficult to sort a single colony of RP. Nevertheless, single red colony of the intratumoral RP (i-RP) was carefully picked up and cultured for further experiments. i-RP was verified as an association of RP and PM by colony assays, fluorescent microscopy, and gene identifications. The ratio of microbial consortium was perpetually RP:PM = 97:3 even after repetitive subculturing as confirmed using fluorescence microscopy and colony assays.

Moreover, super-purified RP (sp-RP) was successfully obtained from PM-swarmed single red colony of i-RP using a fine-tipped syringe needle (∼ 400 μm in diameter) under stereo microscopy. sp-RP was identified to be 100% pure RP using genetic tests (**Supplementary Table S2**).

### Antitumor efficacy of functional bacteria

*In vivo* antitumor therapeutic efficacy of isolated functional bacteria was investigated using a murine Colon-26 carcinoma tumor syngeneic model (**Figure 2A**). Tumor volumes were measured for 120 days after starting the therapeutic regime. Colon-26-bearing immunocompetent mice were intravenously (i.v.) injected with each bacterial suspension or PBS.

**Figure 2.**
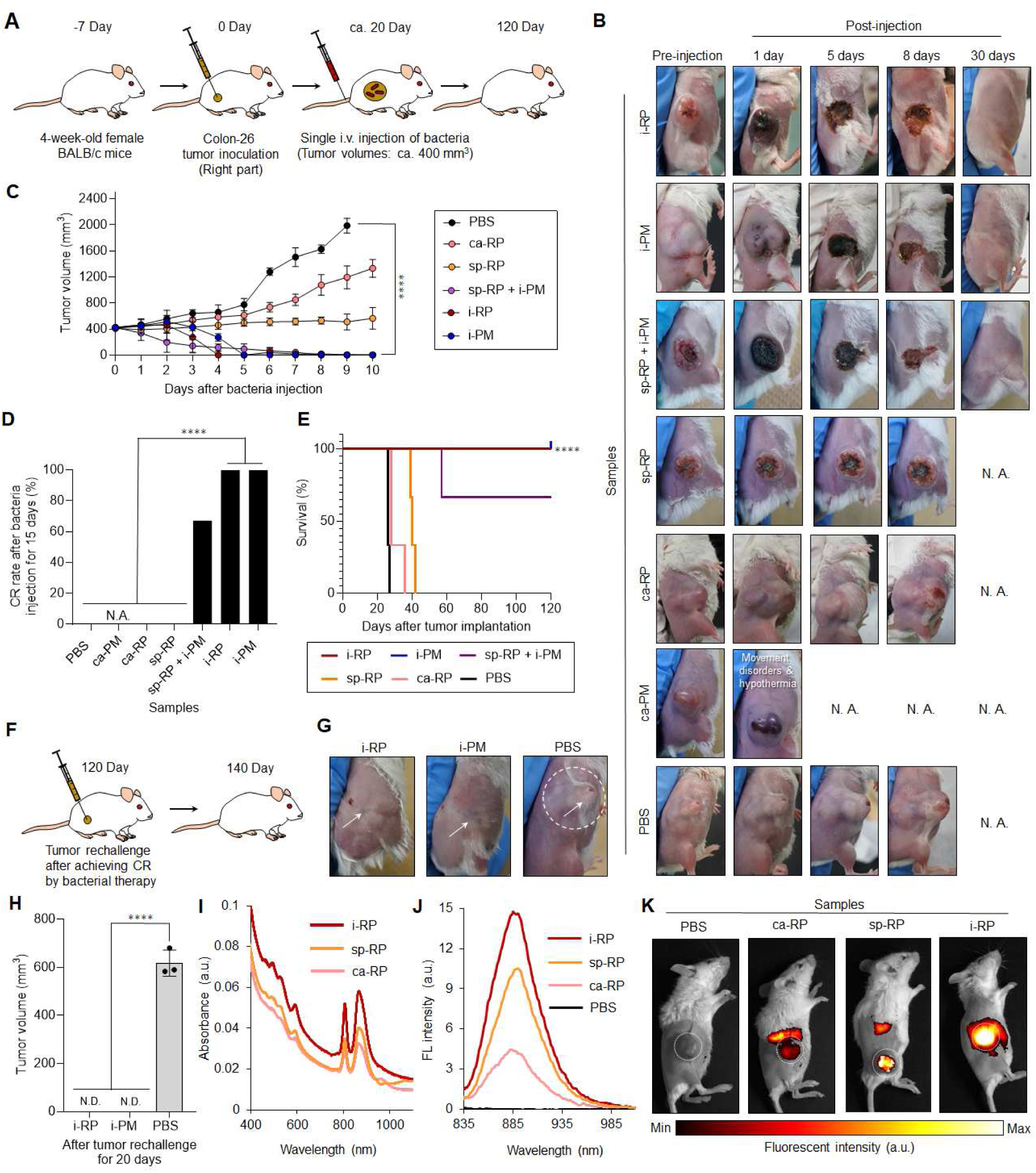
Bacteria-based theranostics for Colon-26-tumor-bearing mice. (**A**) Schematic illustration of *in vivo* Colon-26 carcinoma antitumor tests using various functional bacteria. (**B**) Images of mice after each treatment. N. A., not available. (**C**) *In vivo* anticancer effect of functional bacteria. The PBS or suspension of bacteria was intravenously injected into Colon-26-bearing mice. Data are represented as mean ± standard errors of the mean (SEM); n = 3 biologically independent mice. ****, *p* < 0.0001. (**D**) Complete response (CR) rate of Colon-26-tumor-bearing mice (n = 3 biologically independent mice) after bacteria or PBS injection after 15 days. ****, *p* < 0.0001. (**E**) Kaplan–Meier survival curves of Colon-26-tumor-bearing mice (n = 3 biologically independent mice) after tumor implantation for 40 days. Statistical significance was calculated by comparison with the PBS group. ****, *p* < 0.0001. (**F**) Schematic illustration of tumor rechallenge study after bacterial cancer therapy. Second Colon-26 tumor inoculation into mice (right side) after bacterial treatments or the control experiment with PBS injection are performed at day 120. (**G**) Images of mice in each group after second tumor inoculations assessed after 20 days. The white arrows represent the position of tumor inoculation. The white dashed circle displays the location of solid tumor. (**H**) The tumor volumes for each treated mice (n = 3 biologically independent mice) after tumor inoculation after 20 days. N. D., not detectable; ****, *p* < 0.0001. (**I**) UV‒Vis‒NIR absorbance of various RP suspensions. (**J**) Fluorescent emission spectra of various RP suspensions excited at 805 nm. (**K**) *In vivo* NIR fluorescent bio-imaging of Colon-26-tumor-bearing mice after intravenous injection of PBS (200 µL) or each RP suspension (ca-RP, sp-RP, and i-RP) (200 µL, 1 × 10^9^ CFU) for 5 days. White dashed circles represent the location of tumors.

PBS injections did not influence tumor growth. Meanwhile, to our astonishment, i-RP, i-PM, and sp-RP + i-PM achieved dramatic tumor suppression and complete response (CR) after a single-dose administration (**Figure 2B‒2D**). Apparently, tumors completely disappeared after i.v. injection of i-RP, i-PM, or sp-RP + i-PM for 30 days, and further provided an excellent prognosis with no reoccurrence after treatment. CR rates of i-RP and i-PM were higher than that of sp-RP + i-PM indeed because tumor recurrence was observed in the sp-RP + i-PM group after bacterial injections for 10 days. For accomplishing CR of tumors, optimal bacterial counts of i-RP and i-PM were found to be 1 × 10^9^ CFU and 1 × 10^8^ CFU, respectively (**Supplementary Figure S1**).

Although other bacteria such as sp-RP and ca-RP showed partial response of tumor, these groups were not as effective as i-RP, i-PM, and sp-RP + i-PM. This suggests that the microbial consortium of RP and PM possesses anticancer efficacy in comparison with pure bacteria (sp-RP and ca-RP) presumably because such association can lead to synergetic benefits due to bacterial cross-talks such as biochemical reactions and interbacterial signalling^23, 24^.

Different strains of PM such as PM Hauser (ATCC 35659) are reported to possess oncolytic effects in murine tumors^25^. However, the condition of mice deteriorated a day after administration with ca-PM (NBRC 3849; the same strain as ATCC 35659). Movement disorders (crouching position and shivering) and hypothermia were observed in the ca-PM injected mice, hence further *in vivo* tests using ca-PM were immediately terminated. This result indicated that i-PM could be attenuated while keeping strong anticancer efficacy because biocompatible i-PM was obtained from living mice. Herein, this is the first report showing that isolated intratumoral bacteria demonstrate dramatic CR of tumor after a single administration of bacteria.

The survival rate of mice after i.v. injection of i-RP or i-PM was significantly prolonged compared to the control groups (sp-RP, i-RP, and PBS) (*p* < 0.0001), showing a 100% survival after 120 days (**Figure 2E**). No significant weight loss was observed in any mice, indicating that bacteria exhibit low systemic toxicity except for ca-PM (**Supplementary Figure S2**).

To investigate the effect of bacterial treatment on immunological memory response *in vivo*, we used a tumor rechallenge model (tumor-free mice surviving from a previous survival rate test and naive mice). Tumor developments after first inoculation were conducted for CR-achieved mice by treatment with i-RP (the microbial consortium of sp-RP and i-PM) or i-PM at day 120 (**Figure 2F**). Interestingly, substantial antitumor immunity in the Colon-26 tumor model could be acquired by bacterial treatments probably because of recruitment of immune cells in the tumor microenvironment^26, 27^. In effect, 100% of mice treated with i-RP or i-PM successfully rejected tumor rechallenge (via subcutaneous injection of 1 × 10^6^ Colon-26 cells), demonstrating that functional bacteria can induce durable antitumor immunity (**Figure 2G and 2H**). More surprisingly, these treated mice were healthy without any tumor recurrence for more than 300 days in the whole experimental scheme even after tumor rechallenge. Besides, tumor rechallenge study of the PBS-injected control mice revealed massive tumor formation. CD4^+^ and CD8^+^ memory T cells were found to be increased in the i-PM and i-RP treated tumors; hence, we hypothesised that an inhibitory effect on tumor growth could be induced by these memory T cells (**Supplementary Figure S3**)^27^. Therefore, anticancer strategy using intratumoral bacterial and its microbial consortium could effectively kill primary tumors in response to injected external bacteria, and elicit immunological memory by stimulating systemic antitumor immunity. The intratumoral bacterial platform proposed in this study might offer a new strategy for the improvement of the therapeutic outcomes and the inhibition of tumor recurrence. These data clearly suggest that functional bacteria could be useful for long-term tumor-specific protection.

We previously reported that ca-RP suspensions have specific absorbance in deep tissue-penetrable NIR region at around 805 nm and 850 nm derived from light-harvesting-1-reaction centre (LH1-RC) and peripheral light-harvesting-2 (LH2) antenna protein nanocomplexes^14, 28^. The various RP solutions (i-RP, sp-RP, and ca-RP) prepared in this study displayed the characteristic NIR absorbance (**Figure 2I**) and fluorescence (FL) by NIR excitation at 805 nm (**Figure 2J**). Among them, i-RP was effectively capable to absorb NIR light and expressed strongest NIR FL. In addition, *in vivo* bio-imaging revealed significantly increased FL in all RP strains in targeted tumors (**Figure 2K**). Among them, NIR FL derived from tumor specificity of i-RP was superior than ca-RP and sp-RP. It is known that the tumor-isolated strains present in a tumor growing in mice allow increased targeting of the tumor cells by the immune system^29, 30^. Therefore, these strong optical properties of i-RP might be attributed to the unique biological screening of tumor-specific mutants obtained from a solid tumor surviving in a harsh environment compared to the external RP, resulting in excitometabolism of LH1-RC and LH2 nanocomplexes. Indeed, microbial consortium of i-RP might also improve the optical properties because bacterial communications are known to facilitate metabolism^23, 24^. These results indicated that i-RP could serve as a useful NIR FL probe in mice, and would also be effective as a therapeutic agent for multidimensional tumor targeting and elimination.

### Biological distribution of functional bacteria in tumor

Biological distribution of functional bacteria in cancer cells, spheroids, and tumor tissues were investigated to clarify the tumor penetration and tumor eradication capacity of the i-RP and i-PM (**Figure 3**). Fluorescent microscopy was used for convenient visualisation of these properties of bacteria. Although i-RP has naturally strong NIR FL derived from LH1-RC and LH2 nanocomplexes (**Figure 3A**), i-PM does not have FL in any regions to explore their properties by fluorescent microscopy (**Supplementary Figure S4**). Thus, nanoengineering method using indocyanine green-encapsulating Cremophor® EL (ICG‒CRE) nanoparticle was firstly applied to donate NIR FL to i-PM (**Figure 3B**)^31^. Synthesised ICG‒CRE modified i-PM (ICG‒CRE‒i-PM) exhibited unique optical absorbance and fluorescence properties (**Supplementary Figure S5**). Viability of i-PM could be kept more than 90% during functionalisation with ICG‒CRE nanoparticles. The content of the loading and the internalisation of ICG was approximately 140 µg/mL for 2.0 × 10^9^ CFU/mL of i-PM.

**Figure 3.**
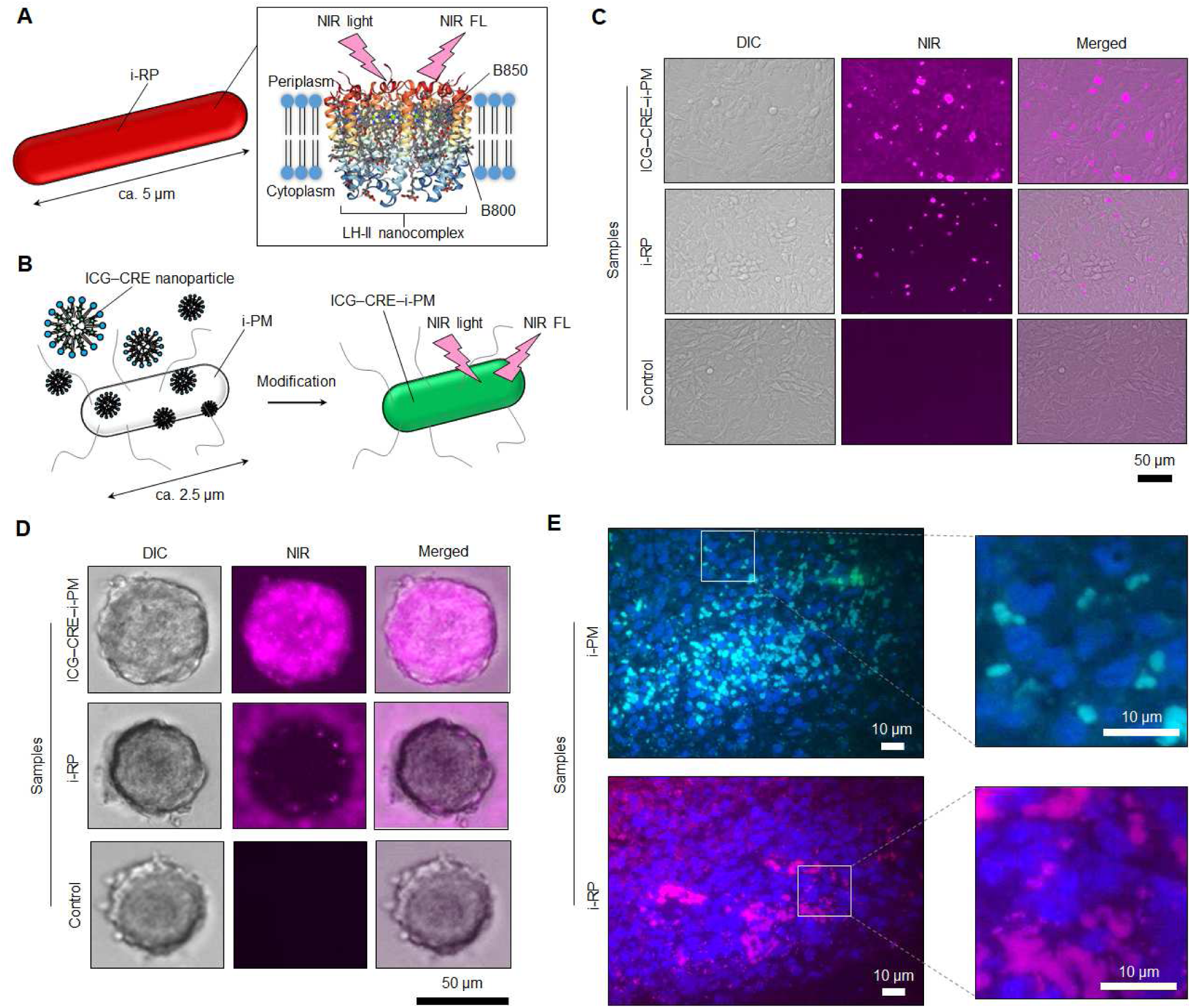
*In vitro* and *in vivo* biological distribution of bacteria in cancer cells, spheroids, and tissues. (**A**) Schematic illustration of NIR light-triggered NIR fluorescence (FL) detection of i-RP. (**B**) Scheme of NIR fluorescent ICG‒CRE‒i-PM synthesis. (**C**) FL images of live Colon-26 cells after treatment with ICG‒CRE‒i-PM and i-RP for 4 h at 37 °C. The pink FL is from bacteria. (**D**) Fluorescent imaging of cancer spheroids incubated with ICG‒CRE‒i-PM and i-RP for 24 h for evaluation of tumor penetration ability. (**E**) Observation of bacterial distribution in a solid tumor tissue by FISH analysis. Bacterial cells and colonies of i-PM and i-RP are light blue and pink in colour, respectively. Cancer cells (blue) was counterstained with 4’,6-diamidino-2-phenylindole (DAPI).

Next, intracellular distribution of i-RP and ICG‒CRE‒i-PM in Colon-26 cells were evaluated using fluorescent microscopy which particularly revealed intracellular ICG‒CRE‒i-PM uptake in cells after incubation at 37 °C for 4 h (**Figure 3C** and **Supplementary Figure S6**). Here, NIR FL of ICG‒CRE‒ i-PM (pink coloured dots) revealed its uniform distribution in the cell cytoplasm. Though i-RP showed NIR FL in cells after incubation at the same condition, its intensity was not stronger than that of the cells harbouring ICG‒CRE‒i-PM. Moreover, i-PM is highly motile peritrichously flagellated microbe, and average length of i-PM (∼ 2.5 µm) is smaller than that of i-RP (∼ 5 µm), resulting in effective internalisation of i-PM into cells. The control cells without bacterial treatment did not show any FL. Meanwhile, after incubation of cells with both bacteria (i-RP and ICG‒CRE‒i-PM) at 4 °C for 4 h, NIR FL assessed from bacteria was not strong, suggesting that the bacterial internalisation into cells was energy dominant^32^. Herein, endocytosis, size and shape of bacteria, and flagella-driven motility are important factors for potential intracellular migrations of functional bacteria.

We further studied the tumor penetration capacity of the ICG‒CRE‒i-PM and i-RP in a spheroid model that originated from Colon-26 cancer cells (**Figure 3D**). Compact sized and highly motile ICG‒ CRE‒i-PM could effectively enter into tumor spheroids compared to i-RP. This reveals that the mobility of bacteria aids in penetrating the core region of the solid tumor in an effective way.

However, it is difficult to explain the reason for high tumor specificity of bacteria based on these *in vitro* “static” tests because both hypoxic environments of tumor and dynamic blood flow are principally essential for the cancer selectivity of bacteria. In fact, various anaerobic bacteria can recognise hypoxia microenvironment in a tumor after their systemic circulation in huge blood vessel networks in the body of a mouse. Thus, by using microbial fluorescence in situ hybridisation (FISH) assay^33^, intratumoral distributions of i-PM and i-RP cells and their colonies were observed in tumors after i.v. injections of i-PM and i-RP through the tail vein of Colon-26-bearing mice (**Figure 3E**). The control (PBS injection) revealed that the native tumor-resident PMs (green coloured dots) were also present in a tumor, whereas any red fluorescent derived from RP were not observed at all (**Supplementary Figure S7**). Herein, we conclude that both i-PM and i-RP are able to deeply penetrate a solid tumor with the help of blood flow and anaerobically proliferate in tumor hypoxic conditions.

### Mechanism of tumor suppression by functional bacteria

To investigate the immunological mechanism regarding solid tumor suppression by bacterial injection, hematoxylin and eosin (H&E) and immunohistochemical (IHC) staining analyses were performed (**Figure 4A** and **Supplementary Figure S8**). IHC staining of F4/80 (macrophage marker), CXCR4 (neutrophil marker), NKp46 [natural killer (NK) cell marker], CD3 (T cell marker), and CD19 (B cell marker) were observed to understand the immunological reactions for the regulation of tumor suppression by functional bacteria. i-PM apparently exhibited the expression of all immunological biomarkers. This indicates that total mobilisation of immune cells are activated in the tumor milieu by i.v. injection of i-PM. Another highly antitumor bacteria i-RP and its mixture (sp-RP + i-PM) significantly stimulate macrophage, and T and B cells those are also aggressive immunological cells for tumor elimination. Other control bacterial groups of ca-RP and sp-RP were not effective for immunological activation rather than i-PM and i-RP. PBS injection did not influence on expression of immunological biomarkers.

**Figure 4.**
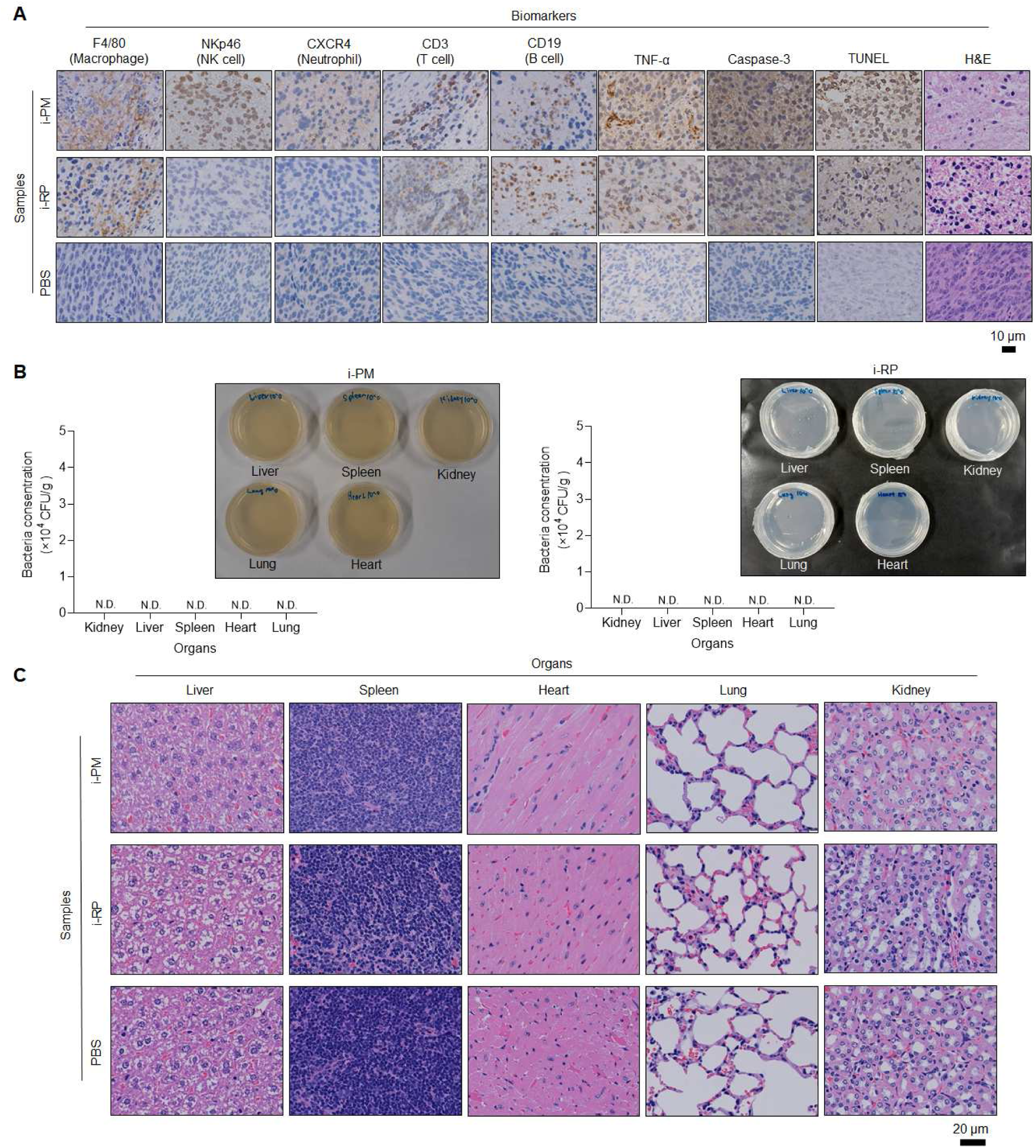
Mechanism of tumor suppression by functional bacteria. (**A**) IHC (F4/80, NKp46, CXCR4, CD3, CD19, TNF-α and caspase-3), TUNEL, and hematoxylin and eosin (H&E) stained tumor tissues collected from different groups of mice at day 1 after respective treatments (PBS, i-RP and i-PM). (**B**) Numbers and images of bacterial colony of organs in Colon-26-tumor-bearing mice after i.v. injection of i-PM (left) (200 µL, 1 × 10^9^ CFU) or i-RP (right) (200 µL, 1 × 10^9^ CFU) after 30 days. Data are represented as mean ± SEM; n = 3 independent experiments. N. D., not detectable. (**C**) H&E staining in conventional organs sectioned after i.v. injection of i-PM (left) (200 µL, 1 × 10^9^ CFU) or i-RP (right) (200 µL, 1 × 10^9^ CFU) after 30 days.

In addition, tumor necrosis factor-α (TNF-α), which is a multifunctional cytokine secreted primarily by macrophages, was particularly expressed in i-PM group although other bacterial groups indicate sufficient TNF-α expression.

Caspase-3 staining assay represents many apoptotic cells especially in i-PM, i-RP, and sp-RP + i-PM groups. Obvious caspase-3 staining in tumor slices after treatments with these intratumoral functional bacteria and microbial consortium indicates massive apoptotic cell death evoked by immunological stimulation features from these bacteria. Simultaneously, terminal deoxynucleotidyl transferase (TdT)-mediated 2’-deoxyuridine, 5’-triphosphate (dUTP) nick end labelling (TUNEL) assay was also commanded to further support strong *in vivo* anticancer mechanism for the intratumoral bacteria. Treatment of intratumoral bacteria (i-PM, i-RP, and sp-RP + i-PM) resulted in enhanced cancer cell apoptosis, as indicated by increased TUNEL-positive cells. Other control bacterial strains (ca-RP and sp-RP) somewhat contribute on apoptotic TUNEL colour development, whereas the control group of PBS alone did not exhibit apoptotic TUNEL colour development within the tumor mass.

Functional bacteria i-PM, i-RP, and sp-RP + i-PM groups demonstrated obvious tumor damage with intercellular fragmentation by H&E staining assay, further revealing antitumor effectiveness. Although other control bacterial injection groups (ca-RP and sp-RP) also showed tumor degeneration, the antitumor therapeutic effectiveness was lower than that observed in the intratumoral bacteria i-PM and its microbial consortium of i-RP and sp-RP + i-PM groups. On the other hand, healthy pathological features, such as tight arrangement and nuclear atypia, were observed in control PBS alone group.

No matter what functional bacteria i-PM and i-RP (microbial consortium of sp-RP and i-PM) can evoke aggressive immune responses in tumor microenvironment, we believe that it would not undergo immunological crisis in other healthy organs. In fact, these bacterial colonies were not detectable post-injection at day 30 in vital organs, such as kidney, liver, spleen, heart, and lung, after treatments in Colon-26 bearing mice using colony counting assays (**Figure 4B**). It clearly indicates that functional bacteria are simultaneously and naturally disappeared after tumor elimination so that it is not necessary to prescribe an antibiotic.

Furthermore, functional bacteria i-PM and i-RP did not show any toxicity in tissues after i.v. injection for 30 days (**Figure 4C**). H&E staining analyses demonstrated that the tissues of post i.v. injection of i-PM and i-RP totally resemble that of control (PBS buffer).

The complete blood counts (CBCs) and blood biochemical parameters of mice were also evaluated after i.v. injecting PBS or bacterial suspension containing i-PM or i-RP for 30 days (**Supplementary Tables S3 and S4**). No significant difference was observed between bacterial administration and control PBS-injected mice groups. Herein, we concluded that the bacterial suspensions were not toxic for *in vivo* administrations.

### Therapeutic extensibility of functional bacteria against various types of cancer

Our bacterial cancer immunotherapies were designed to work in conjunction with immune system to increase native antitumor responses. The functional intratumoral bacteria and its microbial consortium were effective against immunogenic Colon-26 malignant tumor. Finally, presence of such functional bacteria was investigated in treatments of other types of cancer which have different immunogenicity (**Figure 5**).

**Figure 5.**
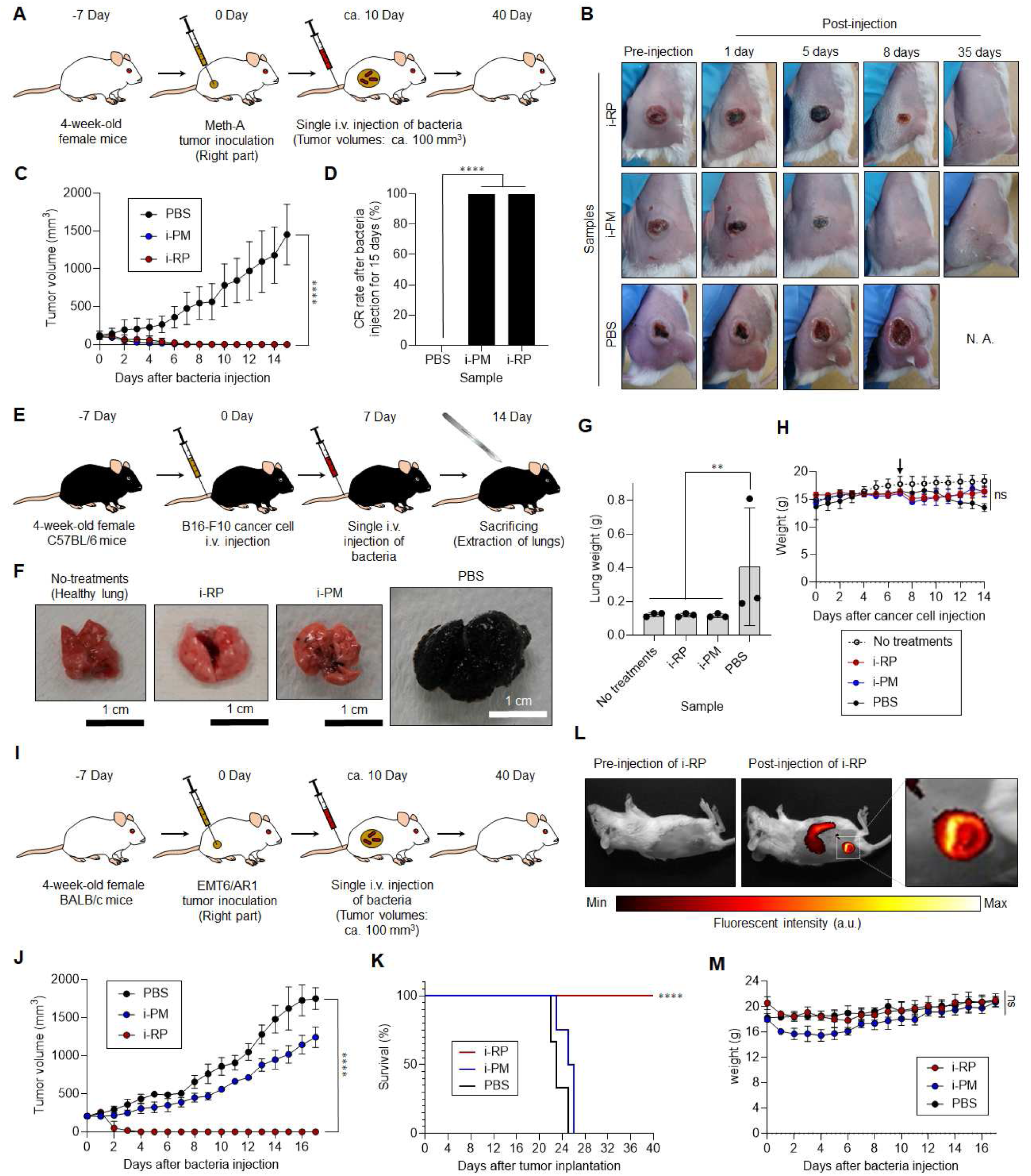
Antitumor efficacy of functional bacteria against various types of tumor-bearing mice. (**A**) Schematic illustration of *in vivo* sarcoma antitumor tests using functional bacteria. (**B**) Images of mice after each treatment. (**C**) *In vivo* anticancer effect of functional bacteria. The PBS or suspension of bacteria was i.v. injected into Meth-A-bearing mice. Data are represented as mean ± SEM; n = 3 biologically independent mice. ****, *p* < 0.0001. (**D**) CR rate of sarcoma-bearing mice (n = 3 biologically independent mice) injected with bacteria or PBS assessed after 15 days. ****, *p* < 0.0001. (**E**) Schematic illustration of *in vivo* antitumor tests using functional bacteria against metastatic lung cancer model. (**F**) Images of lungs after each treatment. (**G**) Weight of lungs after each treatment. Data are represented as mean ± SEM; n = 3 independent experiments. **, *p* < 0.01. (**H**) Average body weight of the mice after bacterial and control treatments during the experimental duration. Black arrow shows the time point of sample injection (Day 7). Data are represented as the mean ± SEM; n = 3 biologically independent mice. ns, not significant. (**I**) Schematic illustration of *in vivo* antitumor tests using functional bacteria against drug-resistant cancer model. (**J**) Relative volumes of the tumors on the right flank of mice after i.v. injection of i-RP, i-PM, or PBS. Bacterial concentrations of i-RP and i-PM are 12.5 × 10^9^ CFU/mL and 5 × 10^8^ CFU/mL, respectively. Data are presented as mean ± SEM (n = 3 biologically independent tests), ****, *p* < 0.0001 (Student’s t-test used for analysis in comparison to PBS). (**K**) Kaplan-Meier survival curves of EMT6/AR1-bearing mice (n = 3 biologically independent mice) after tumor implantation for 40 days. Statistical significance was calculated in comparison with PBS group. ****, *p* < 0.0001. (**L**) NIR FL images of the mouse before and after i.v. injection of i-RP (5 × 10^9^ CFU/mL) for 3 days. (**M**) Weight of mice after each treatment. Data are represented as means ± SEM; n = 3 independent experiments. ns, not significant.

Both of i-RP and i-PM represented dramatic effectiveness against an immunogenic murine skin sarcoma (Meth-A)-bearing tumor model and successfully accomplished 100% of CR rate of tumors (**Figure 5A–5D**). These bacteria also improved survival rate of Meth-A bearing mice (**Supplementary Figure S9**). Furthermore, mortality rates were 0% after i.v. injections with i-RP or i-PM after 40 days. There were no significant differences on weight of mice between PBS and these bacterial injected groups, demonstrating that the bacteria did not have any side effects on Meth-A bearing mice (**Supplementary Figure S9**).

More interestingly, these bacteria also showed potent antitumor efficacy against metastatic lung cancer derived from non-immunogenic murine melanoma (B16F10) (**Figure 5E–5H**). Lungs excised from no-treatment exhibited normal physiological features, and the lungs of mice treated with i-RP resembled those of controls after i.v. injection (**Figure 5F**). Although i-PM-treated lungs had slightly metastatic pulmonary nodules (small black dots), it looked almost similar to native lungs. Meanwhile, the lungs of B16F10 melanoma-bearing mice revealed enlarged black tumors. The lungs of mice treated with both (B16F10 implantation and i-RP or i-PM i.v. injection) clearly exhibited a significantly reduced tumor weight versus B16F10 melanoma-bearing mice (**Figure 5G**). The average body weights of mice treated with both B16F10 cells and i-RP or i-PM were determined, and there were no significant differences compared to that of the control groups (no-treatment and treatment with B16F10 cells and PBS) (**Figure 5H**).

Moreover, poorly immunogenic and drug-resistant mouse mammary tumor (EMT6/AR1)-bearing mice were also effectively cured by i-RP treatment (**Figure 5I–5M**). In fact, i-RP injection successfully achieved CR of tumors (**Figure 5J and Supplementary Figure S10**). Survival rates of mice were prolonged by the single i.v. administration of i-RP at least 40 days (**Figure 5K**). Optical nanofunction of i-RP was also useful for effective NIR FL marking of tumor location (**Figure 5L**). Meanwhile, i-PM represented a mild antitumor efficacy against the extensive drug-resistant model (**Figure 5J**). Both bacteria were totally safe for EMT6/AR1-bearing mice as well as in other cancer models, resulting in no adverse effect on body weight of the mice (**Figure 5M**). These results demonstrate that immunological efficacies and optical property of functional bacteria are highly effective for eradication of drug-resistant aggressive cancers.

Finally, anticancer efficacies of other intratumoral bacteria [*Lactococcus sp.* (i-LS) and *Enterococcus faecalis* (i-EF)], which were isolated from solid tumor biopsies without i.v. injection of RP, were further investigated using Colon-26-bearing mice (**Supplementary Figure S11**). Isolations of i-LS and i-EF were successfully assessed using similar means of extractions of i-RP, i-PM, and sp-RP except the use of Luria broth (LB) medium (**Supplementary Figure S11A**). Interestingly, homology of i-LS was closest to *Lactococcus formosensis* (97.69%) or *Lactococcus garvieae* (97.48%) by Basic Local Alignment Search Tool (BLAST) analyses. In other words, gene identification exhibit that i-LS is potentially a new strain because the similarity of 16S rRNA gene of i-LS is less than 98.7% in comparison with the data-based bacterial strains (**Supplementary Table S5**)^34^. Meanwhile, i-EF was confirmed as 100% pure EF using gene identification (**Supplementary Table S6**). In any case, both i-LS and i-EF showed tumor suppression effect in Colon-26-bearing mice although slight body weight reductions were observed after their injections (**Supplementary Figure S11B‒11F**). i-LS represented stronger anticancer efficacy and achieved 33% CR. These results clearly indicate a high potentiality of various intratumoral microbiota as a powerful antitumor medicine. Therefore, we concluded that tumor-resident functional bacteria obtained could serve as a universal immuno-activatable therapeutic “living” agent in various syngeneic cancer models.

## Discussion

Earlier studies on the correlations between intratumoral bacteria and cancer have mainly concentrated on studying the role of bacteria in tumor pathogenesis and the contribution of metabolism of bacteria to the adverse effects of chemotherapy. However, therapeutic activities of intratumoral bacteria themselves has not been explored. Here, we demonstrate that isolated native intratumoral bacteria and its microbial consortium that can potentially modulate various immunological activities to dramatically eliminate various tumors with or without tumor immunogenicity. They exhibit high biocompatibility in mice as well as antitumor effectiveness with excellent tumor selectivity through single administration. Association of intratumoral bacteria and photosynthetic bacteria can particularly work as not only a potent therapeutic agent but also as a beneficial cancer diagnosis probe, indicating highly tumor-specific strong NIR FL emission and immunological elimination under tissue-penetrable NIR light.

Number of cancer patients in the world has drastically increased, resulting in annual spending on anticancer drugs more than US$ 100 billion, and is predicted to rise continually^35^. However, despite the high demand, the price of advanced anticancer medicine has skyrocketed. Moreover, at the peak of the COVID-19 pandemic, tens of millions of people globally lost their jobs and health coverage^36^. Such unfortunate situation affected groups suffering from both cancers and poverty. To that end, the functional bacteria discovered in this study can be abundantly proliferated in inexpensive media without any complicated process, indicating that they may resolve the troublesome cost issue of anticancer medicines. Further, these functional bacteria would be a useful “hardware” in which genes and microbial surface can be genetically and chemically manipulated in combination with emerging genetic and nano engineering approaches, for obtaining higher therapeutic performances and multifunction by modulating metabolic pathways and/or loading with nanoparticles. The bacterial platform technology, that does not require substantial petrochemicals, may also shed light on global environmental concerns because functional bacteria could be alternative ecofriendly medicinal materials instead of traditional organic anticancer drugs typically synthesised from fossil fuels with massive carbon dioxide emissions^37^.

More importantly, this study is the first to draw attention to potentially myriad intratumoral bacteria, also called microbial dark matter, as an independent natural active anticancer medicine for future innovative cancer treatment. We envision that tailor-made intratumoral microbiota derived from individual cancer patients might have unique anticancer efficacy as a personalised medicine. Bacterial cancer therapy is now experiencing renaissance, and the gold rush of research into various intratumoral bacteria for cancer treatment has just begun.

## Materials and Methods

### Bacterial strains and growth

The bacterial strains tested in this study were purchased from the National Institute of Technology and Evaluation Biological Resource Center (NBRC) (Chiba, Japan). Commercially available *Rhodopseudomonas Palustris* (ca-RP) (NBRC 16661) was grown anaerobically in liquid cultures at 26‒30 °C in ATCC 543 medium under tungsten lamps. Commercially available *Proteus mirabilis* (ca-PM) (NBRC 3849) was anaerobically cultured at 30 °C in NBRC 802 medium. All reagents for bacterial culturing were obtained from Nacalai Tesque (Kyoto, Japan), Tokyo Chemical Industry (Tokyo, Japan), and FUJIFILM Wako Pure Chemical (Osaka, Japan).

### Cell culture

Murine colon carcinoma (Colon-26) cells were obtained from the Japanese Collection of Research Bioresources Cell Bank (Tokyo, Japan). Murine skin sarcoma (Meth-A) and murine melanoma (B16F10) cell lines were purchased from Cell Resource Center for Biomedical Research, Institute of Development, Aging and Cancer Tohoku University. Drug-resistant mouse mammary tumor (EMT6/AR1) cell line was obtained from KAC Co., Ltd. (Tokyo, Japan). The Colon-26, Meth-A, and B16F10 cell lines were cultured in Roswell Park Memorial Institute 1640 Medium (Gibco, Grand Island, NY, USA) containing 10% foetal bovine serum, 2 mM L-glutamine, 1 mM sodium pyruvate, gentamycin, and penicillin-streptomycin (100 IU/mL). EM6/AR1 cell line was cultured in cell growth medium No.104 (KAC Co., Ltd.) containing doxorubicin (1 µg/mL). Cells were maintained at 37 °C in a humidified incubator containing 5% CO_2,_ were cryopreserved in multiple vials and stored in liquid nitrogen. Cell stocks were regularly revived to avoid the genetic instabilities associated with high passage numbers.

### Isolations of functional bacteria from solid tumors

The animal experiments were conducted in accordance with the protocols approved by the Institutional Animal Care and Use Committee of JAIST (No. 04-002). Female BALB/cCrSlc mice (n = 8; 4-weeks-old; average weight = 15 g) were obtained from Japan SLC (Hamamatsu, Japan). Mice bearing the Colon-26 cell-derived tumors were generated by injecting 100 μL of the culture medium/Matrigel (Dow Corning, Corning, NY, USA) mixture (v/v = 1:1) containing 1 × 10^6^ cells into the right side of backs of the mice. After approximately 20 days, when the tumor volumes reached ∼400 mm^3^, the mice were intravenously injected with 200 μL culture medium containing ca-RP (5 × 10^9^ CFU/mL). After 2 days, the tumors were carefully excised. After homogenising thoroughly with a homogeniser pestle in 1 mL of PBS solution at 4 °C, the mixture was shaken for 20 min at a speed of 380 rpm/min at 15 °C. The supernatant was diluted 10 times with PBS, and then a sample (100 μL) was inoculated onto an agar plate. After incubation for 3 days under anaerobic conditions and tungsten lamps, the red bacterial colonies derived from *Rhodopseudomonas Palustris* were formed on plates. Several colonies were picked, and cultured anaerobically in ATCC 543 medium at 26‒30 °C under tungsten lamps for 3 days. Grey-coloured bacteria solution was subcultured in conventional ATCC 543 medium or ATCC 543 medium without L-cysteine (Cys). Subcultured red bacterial suspension, that was prepared in ATCC 543 medium without Cys, was disseminated onto an ATCC 543 agar plate with 0.1% sodium deoxycholate (SD) and without Cys. After 7 days, red and white colonies were observed on the plate. Single red colony was carefully picked up using an inoculating loop, and then cultured in an ATCC 543 medium without Cys for 3 days. Prepared red bacterial solution [isolated *Rhodopseudomonas Palustris* (i-RP)] was used for further tests.

Meanwhile, after subculturing, obtained grey-coloured bacterial solution, that was prepared by conventional ATCC 543 medium, was inoculated onto an ordinary ATCC 543 agar plate containing 0.1% SD. White bacterial colonies were formed on the plate, and single white colony was carefully picked up by an inoculating loop, and then anaerobically cultured in conventional ATCC 543 medium at 26‒30 °C under tungsten lamps for 3 days. Prepared grey bacterial solution [isolated *Proteus mirabilis* (i-PM)] was used for further tests.

Besides, single red colony prepared from i-RP solution was carefully picked up using a 27-gauge syringe needle (SS-01T2719S, 0.4 mm diameter, Terumo, Tokyo, Japan) under stereo microscopy (M2-1139-01; AS ONE, Osaka, Japan), and then anaerobically cultured in an ATCC 543 medium without Cys at 26‒30 °C under tungsten lamps for 3 days. Prepared red solution [super-purified *Rhodopseudomonas Palustris* (sp-RP)] was used for further tests.

The i-RP was identified as microbial consortium of *Rhodopseudomonas Palustris* and *Proteus mirabilis* by various investigations such as colony assay, optical microscopy, and gene identifications with Basic Local Alignment Search Tool (BLAST) analyses using 16S rRNA primers (listed in **Supplementary Table S7**). The gene identifications were performed by BEX Co. (Tokyo, Japan). The ratio of the microbial consortium of i-RP was measured as RP:PM = 97:3 by fluorescent microscopy and colony assay. The i-PM and sp-RP were confirmed as pure bacteria by colony assays, optical microscopy, and gene identifications.

*Lactococcus sp.* (i-LS) and *Enterococcus faecalis* (i-EF) were also isolated from solid tumors (∼400 mm^3^) in Colon-26-bearing mice using syringe needle and stereomicroscopy without i.v. injections of RP, and anaerobically cultured in Luria Broth (LB) agar plate and LB medium at 26‒30 °C under tungsten lamps. i-LS was subcultured in Pearl Core® E-MC64 medium (Eiken Chemical, Tokyo, Japan) at 32 °C using incubator (i-CUBE FCI-280HG; AS ONE). i-EF was subcultured in LB medium (Nacalai Tesque) at 26‒30 °C under tungsten lamps. i-LS and i-EF obtained were genetically identified as pure strains by 16S rRNA techniques and BLAST analyses in similar way to the identifications of other bacterial strains in this study.

### Preparation of ICG‒CRE‒i-PM

Preparation of ICG‒CRE‒i-PM was as reported previously^31^. Briefly, indocyanine green (ICG, 1 mg/mL; Tokyo Chemical Industry) was dissolved in PBS with 5% cremophor EL (Nacalai Tesque) using sonication to obtain ICG‒CRE nanoparticle. The medium containing i-PM was centrifuged at 3,000 × *g* for 5 min at a temperature of 4‒15 °C and washed with PBS. Following this, the concentration of the bacterial suspension was adjusted to 2 × 10^9^ CFU/mL and centrifuged to obtain a bacterial pellet. Then ICG‒CRE (ICG concentration = 1 mg/mL) was added to it and the cell suspension was incubated overnight at 37 °C. Further, the samples were centrifuged to remove the unattached dye molecules, and the modified bacteria were resuspended in PBS. The viabilities of ICG‒ CRE‒i-PM were measured using a bacterial counter (DU-AA01NP-H; PHC Co., Ltd., Tokyo, Japan) in addition to active colony assay.

### Optical characterisations of bacteria

Absorption spectra of bacterial solutions were recorded at 20 °C using a UV–Vis–NIR spectrophotometer (V-730 BIO; Jasco, Tokyo, Japan). Fluorescence of bacterial suspensions were measured using fluorescence spectrometer (FP-8600 NIR Spectrofluorometer; Jasco, Tokyo, Japan). Amount of loaded and internalised ICG into bacteria was calculated from collected supernatant of washed i-PM after modification of ICG‒CRE by a UV–Vis–NIR spectrophotometer.

### FL microscopy imaging of bacteria

Bacterial solutions (20 μL, 5 × 10^8^ CFU/mL) were plated on glass coverslips (AGC Techno glass, Shizuoka, Japan) and then observed using a fluorescence microscopy system (IX73) and cellSens V3.1 software (Olympus) equipped with a mirror unit (IRDYE800-33LP-A-U01; Semrock, Lake Forest, IL, USA) and objectives (×60 magnification, aperture 1.35; UPLSAPO60X, Olympus or ×100 magnification, aperture 0.95; PLFLN100X, Olympus) at room temperature.

### Intracellular penetration of bacteria

Colon-26 cells (2.5 × 10^5^ cells/well) were seeded in poly-L-lysine coated glass bottom dishes (Matsunami glass, Osaka, Japan) and allowed to adhere overnight. Cells were then exposed to 1 × 10^7^ CFU of i-RP or 1 × 10^7^ CFU of ICG‒CRE‒i-PM for 4 h at 4 °C or 37 °C in a fridge or a humidified incubator containing 5% CO_2_. After washing thoroughly with fresh PBS solution, Colon-26 cells were observed using a fluorescence microscopy system (IX73) equipped with a mirror unit (IRDYE800-33LP-A-U01; Semrock, Lake Forest, IL, USA) and an objective (×20 magnification, aperture 0.75; UPLSAPO20X, Olympus) at room temperature.

### Tumor spheroids

Colon-26 cells (1 × 10^4^ cells/well) were seeded in a 3D culture spheroid plate (Cell-able® BP-96-R800; Toyo Gosei, Tokyo, Japan) according to the manufacturer’s instructions provided with the plate. Cells were cultured for 7 days at 37 °C in a humidified incubator containing 5% CO_2_. Medium was replaced every 2 days. Prepared spheroids were then exposed to 1 × 10^6^ CFU of i-RP or 1 × 10^6^ CFU of ICG‒CRE‒i-PM for 24 h at 37 °C in a humidified incubator containing 5% CO_2_. After washing thoroughly with fresh PBS solution, spheroids were observed using a fluorescence microscopy system (IX73) equipped with a mirror unit (IRDYE800-33LP-A-U01; Semrock) and an objective (×20 magnification, aperture 0.75; UPLSAPO20X, Olympus) at room temperature.

### FISH assay

Microbial FISH assay of i-RP and i-PM in a tumor slice was assessed using a *Rhodopseudomonas* (Probe name, Rhodopseud; Accession no. pB-1634; Chromosome Science Labo, Hokkaido, Japan) and *Proteus mirabilis* (Probe name, EUB338; Chromosome Science Labo) specified microbial FISH probes according to the manufacturer’s instructions provided with the kit. Rhodopseud and EUB338 Probes were labelled with cyanine 3 (Cy3) and fluorescein isothiocyanate (FITC), respectively. The specificity of the probes was confirmed by database at ProbeBase (University of Vienna, Vienna, Austria) and the previous literatures^38^. Briefly, after deparaffinising the tumor sections and treating with 0.02% pepsin (FUJIFILM Wako Pure Chemical) /0.1N HCl solution (FUJIFILM Wako Pure Chemical), the sections were stained with Rhodopseud probe (5’-GACTTAGAAACCCGCCTACG-3’) (listed in **Supplementary Table S8**) or EUB338 (5’-GCCCCTGCTTTGGTC-3’) probe (listed in **Supplementary Table S8**) and DAPI (Dojindo Laboratories), and examined using fluorescent microscopy (IX73).

### *In vivo* anticancer therapy

The Colon-26 tumor-bearing mice (female, about 8 weeks; n = 4; average weight = 18 g; average tumor size ∼ 400 mm^3^; BALB/cCrSIc; Japan SLC) were intravenously injected in the tail vein with culture medium (200 μL) containing i-RP (5 × 10^9^ CFU/mL), i-PM (5 × 10^8^ CFU/mL), sp-RP + i-PM (5 × 10^9^ CFU/mL, sp-RP : i-PM = 97 : 3), sp-RP (5 × 10^9^ CFU/mL), ca-RP (5 × 10^9^ CFU/mL), or ca-PM (5 × 10^8^ CFU/mL). The control experiments were also performed using PBS (200 μL). The tumor formation and overall health (viability and body weight) were monitored every day. Further, the tumor volume was calculated using V = L × W^2^/2, where L and W denote the length and width of the tumor, respectively. The survival ratio of Colon-26 tumor-bearing mice (n = 4 biologically independent mice) was also measured during treatments for 40 days. When the tumor volumes reached more than 2,000 mm^3^, the mice were euthanised as the endpoint according to the guidelines of Institutional Animal Care and Use Committee of JAIST.

For studying cancer immunity, the CR-achieved mice (Day 120) by bacterial therapy (n = 3 biologically independent mice) were injected at the right side of backs of mice with 100 μL of the culture medium/Matrigel mixture (v/v = 1:1) containing 1 × 10^6^ cells of Colon-26. After 20 days, no reoccurrences of tumor were carefully assessed during 20 days. The control experiments were also performed after i.v. injections of PBS (200 µL) into BALB/c mice (female, about 8 weeks; n = 3; average weight = 18 g; BALB/cCrSIc; Japan SLC). After 120 days, 100 μL of the culture medium/Matrigel mixture (v/v = 1:1) containing 1 × 10^6^ cells of Colon-26 was subcutaneously injected into the right side of backs of mice. The volumes of tumors were measured after 20 days similar to the aforementioned *in vivo* anticancer therapy against Colon-26 syngeneic mouse model.

For investigating *in vivo* anticancer therapy using a sarcoma model, mice bearing Meth-A cell-derived tumors were generated by injecting 100 μL of the culture medium/Matrigel (Dow Corning, Corning, NY, USA) mixture (v/v = 1:1) containing 1 × 10^6^ cells into the right side of backs of the mice (female, 5 weeks; n = 3; average weight = 18 g; BALB/cCrSIc; Japan SLC). After approximately 10 days, when the tumor volumes reached ∼100 mm^3^, the mice were intravenously injected with 200 μL of culture medium containing i-RP (5 × 10^9^ CFU/mL) or i-PM (5 × 10^8^ CFU/mL). The control experiments were also performed by i.v. injection of PBS (200 μL) into a sarcoma model. The tumor volume, overall health (viability and body weight), and the survival ratio of mice were investigated in the similar way to the anticancer study of Colon-26 tumor-bearing mice.

To obtain a metastatic lung cancer-bearing animal model, female C57BL/6 mice (5 weeks; n = 3; average weight = 17 g; C57BL/6NCrSlc; Japan SLC) were injected with 100 mL of 5 × 10^6^ of murine melanoma (B16F10) cells using a 27-gauge needle through a tail vein (Day 0). 200 μL of culture medium containing i-RP (5 × 10^9^ CFU/mL) or i-PM (5 × 10^8^ CFU/mL) was intravenously injected after B16F10 cell implantation for 7 days. Animals were sacrificed on day 14 and lungs were harvested. In addition, the lung weight of each mouse treated with PBS or bacteria was measured at the same date. The control experiments without any treatments (inoculations of melanoma and bacteria) were also performed.

In order to investigate the efficacy of bacteria against drug-resistant cancer model, 100 μL of the culture medium/Matrigel (Dow Corning, Corning, NY, USA) mixture (v/v = 1:1) containing 1 × 10^6^ of EMT6/AR1 cells were subcutaneously inoculated into the right side of backs of the mice (female, 5 weeks; n = 3; average weight = 18 g; BALB/cCrSIc; Japan SLC). After approximately 10 days, when the tumor volumes reached ∼100 mm^3^, the mice were intravenously injected with 200 μL culture medium containing i-RP (12.5 × 10^9^ CFU/mL) or i-PM (5 × 10^8^ CFU/mL). The control experiments were also performed using PBS (200 μL). The tumor volume, overall health (viability and body weight), and the survival ratio of mice were also studied.

For *in vivo* tumor tests using i-LS and i-EF, the Colon-26 tumor-bearing mice (female, about 8 weeks; n = 3; average weight = 17 g; average tumor size ∼ 100 mm^3^; BALB/cCrSIc; Japan SLC) were intravenously injected in the tail vein with culture medium (200 μL) containing i-LS (5 × 10^9^ CFU/mL) or i-EF (1 × 10^9^ CFU/mL). The control experiments were performed using PBS (200 μL). The tumor volume, overall health (viability and body weight), and the survival ratio of mice were investigated.

### Biodistribution of bacteria in tumor model

The Colon-26 tumor-bearing mice (female, about 8 weeks; n = 3; average weight = 18 g; average tumor size ∼ 400 mm^3^; BALB/cCrSIc; Japan SLC) were intravenously injected in the tail vein with culture medium (200 μL) containing i-RP (5 × 10^9^ CFU/mL) or i-PM (5 × 10^8^ CFU/mL). The control experiments were performed using PBS (200 μL). After 30 days, the organs were carefully excised and weighed. After homogenising thoroughly with a pestle in 1 mL of PBS solution at 4 °C, the mixture was shaken for 20 min at a speed of 380 rpm/min at 15 °C. The supernatant was diluted 10 times with PBS, and a sample (100 μL) was then inoculated onto an agar plate. After incubation under anaerobic conditions for 7 days, the formed bacterial colonies were carefully confirmed. For counting bacterial colonies, the supernatant was diluted 0, 10, 100, and 1000 times with PBS, and then a sample (5 μL) was inoculated onto an agar plate as above-mentioned. Finally, formed bacterial colonies were manually counted.

### *In vivo* fluorescent bio-imaging

In order to monitor the NIR FL intensity caused by the tumor targeting effect of *R. Palustris* (ca-RP, sp-RP, and i-RP) in mice, Colon-26 tumor-bearing mice (female; about 8 weeks; n = 4; average weight = 18 g; average tumor size = 400 mm^3^; BALB/cCrSIc; Japan SLC) were injected intravenously with culture medium containing *R. Palustris* (200 μL, 1 × 10^9^ CFU). The whole body of mice were imaged using an *in vivo* FL imaging system (VISQUE™ InVivo Smart-LF, Vieworks, Anyang, Republic of Korea) with a 3 sec exposure time and indocyanine green (ICG) filter (Ex, 740–790 nm; Em, 810–860 nm) at 5-days post-injection. The FL images were acquired and analysed using CleVue™ software.

### Tumor tissues immunohistochemistry staining

The Colon-26 tumor-bearing mice (female; about 8 weeks; n = 4; average weight = 18 g; average tumor size = 400 mm^3^; BALB/cCrSIc; Japan SLC) were euthanised the next day after administration of PBS (200 µL)/ca-RP (200 µL, 1 × 10^9^ CFU)/sp-RP (200 µL, 1 × 10^9^ CFU)/ sp-RP + i-PM (200 µL, 1 × 10^9^ CFU, sp-RP : i-PM = 97 : 3)/i-PM (200 µL, 1 × 10^8^ CFU)/i-RP (200 µL, 1 × 10^9^ CFU) injection intravenously. Thereafter, the tumor tissues from the different treatment groups were harvested at day 1 for IHC staining. Analysis was performed by Biopathology Institute Co., Ltd. (Oita, Japan) using standard protocols. Briefly, primary tumors were surgically removed, fixed in 10% formalin, processed for paraffin embedding, and then cut into 3–4-μm-thick sections. After incubation with primary antibodies (listed in **Supplementary Table S9**), the sections were stained with hematoxylin and examined using light microscopy (IX73).

### Blood tests

The CBC and biochemical parameters were investigated by Japan SLC and Oriental Yeast Co. (Tokyo, Japan). BALB/cCrSlc mice (female; 10 weeks; n = 5; average weight = 21 g; Japan SLC) were injected in the tail vein with culture medium containing bacteria (200 μL, 1 × 10^6^ CFU) or PBS (200 μL). The blood samples were collected from the inferior vena cava of the mouse after 30 days.

### Statistical analysis

All experiments except for Supplementary Information were performed in triplicate and repeated three or more times. Quantitative values are expressed as the mean ± standard error of the mean (SEM), of at least three independent experiments. Statistical differences were identified using the Student’s *t*-test or one-way/two-way analysis of variance (ANOVA) followed by Tukey tests.

## Author contributions

E. M. led and supervised the project; E. M. conceived and designed the experiments; E. M. prepared the manuscript; Y. G. and S. I. performed the experiments; All the authors analyzed the data, discussed the results and contributed to writing of the manuscript.

## Acknowledgments

This work was financially supported by Japan Society for the Promotion of Science KAKENHI Grant-in-Aid for Challenging Research (Pioneering) (Grant number 22K18440), the Japan Science and Technology Agency for Adaptable and Seamless Technology Transfer Program through Target-driven R&D (Grant Number JPMJTR22U1), Institute for Fermentation, Osaka (IFO), and the Uehara Memorial Foundation. The authors also thank Mr. Satoru Komatsu (Japan Advanced Institute of Science and Technology) and Ms. Sheethal Reghu (Japan Advanced Institute of Science and Technology) for their partially support with some experiments.

## Data availability

All data needed to evaluate the conclusions in the paper are presented in the paper and/or the Supplementary Materials. Additional data related to this paper are available from the corresponding author on reasonable request.

## Supplementary Information

### Supplementary Figures & Tables

**Supplementary Figure S1.**
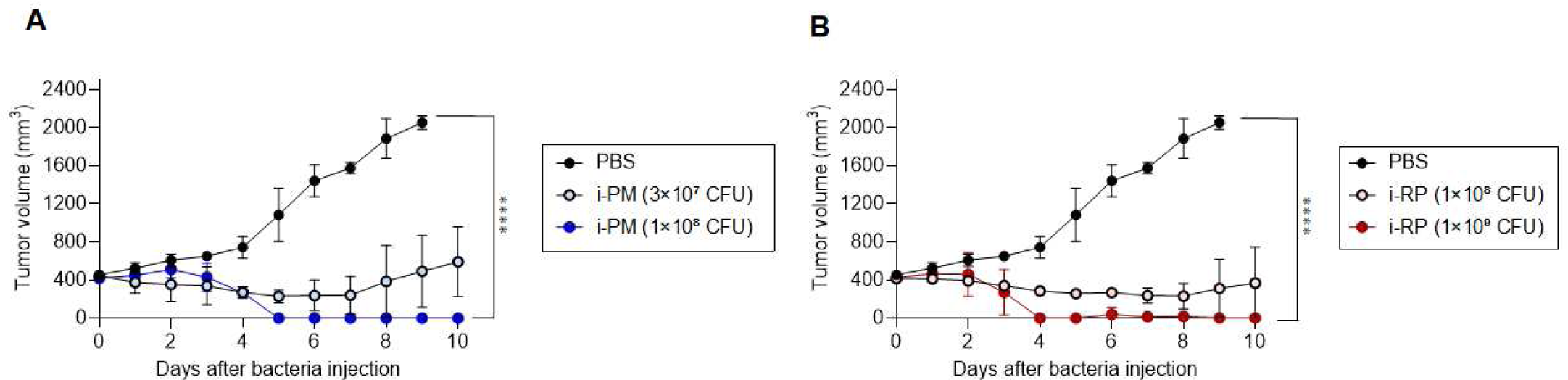
Effect of bacterial concentrations on *in vivo* antitumour efficacy. (**A**) *In vivo* Colon-26 anticancer effect of i-PM at different concentration (3 × 10^7^ CFU and 1 × 10^8^ CFU). Data are represented as mean ± standard errors of the mean (SEM); n = 2 (for 3×10^7^ CFU) or n = 3 (for 1 × 10^8^ CFU and PBS) biologically independent mice. ****, *p* < 0.0001. (**B**) *In vivo* Colon-26 anticancer effect of i-RP at different concentration (1 × 10^8^ CFU and 1 × 10^9^ CFU). Data are represented as mean ± SEM; n = 2 (for 1 × 10^8^ CFU) or n = 3 (for 1 × 10^9^ CFU and PBS) biologically independent mice. ****, *p* < 0.0001.

**Supplementary Figure S2.**
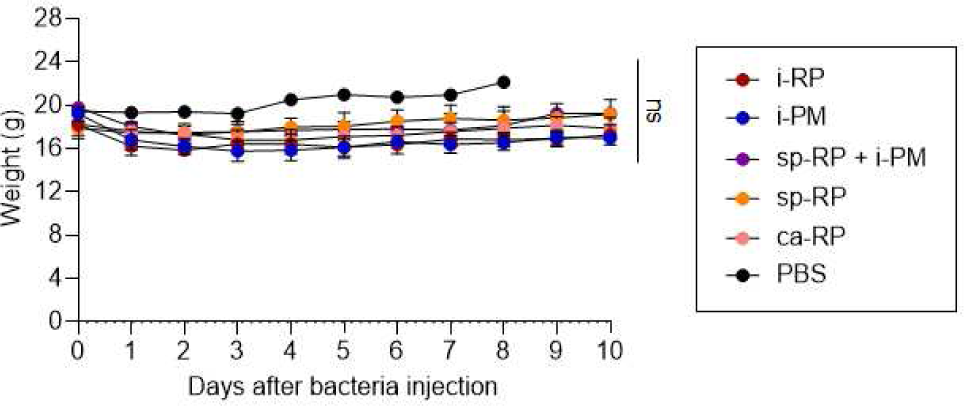
Weight measure after various treatments in Colon-26-tumour-bearing mice. Data are represented as mean ± SEM; n = 3 independent experiments. ns, not significant.

**Supplementary Figure S3.**
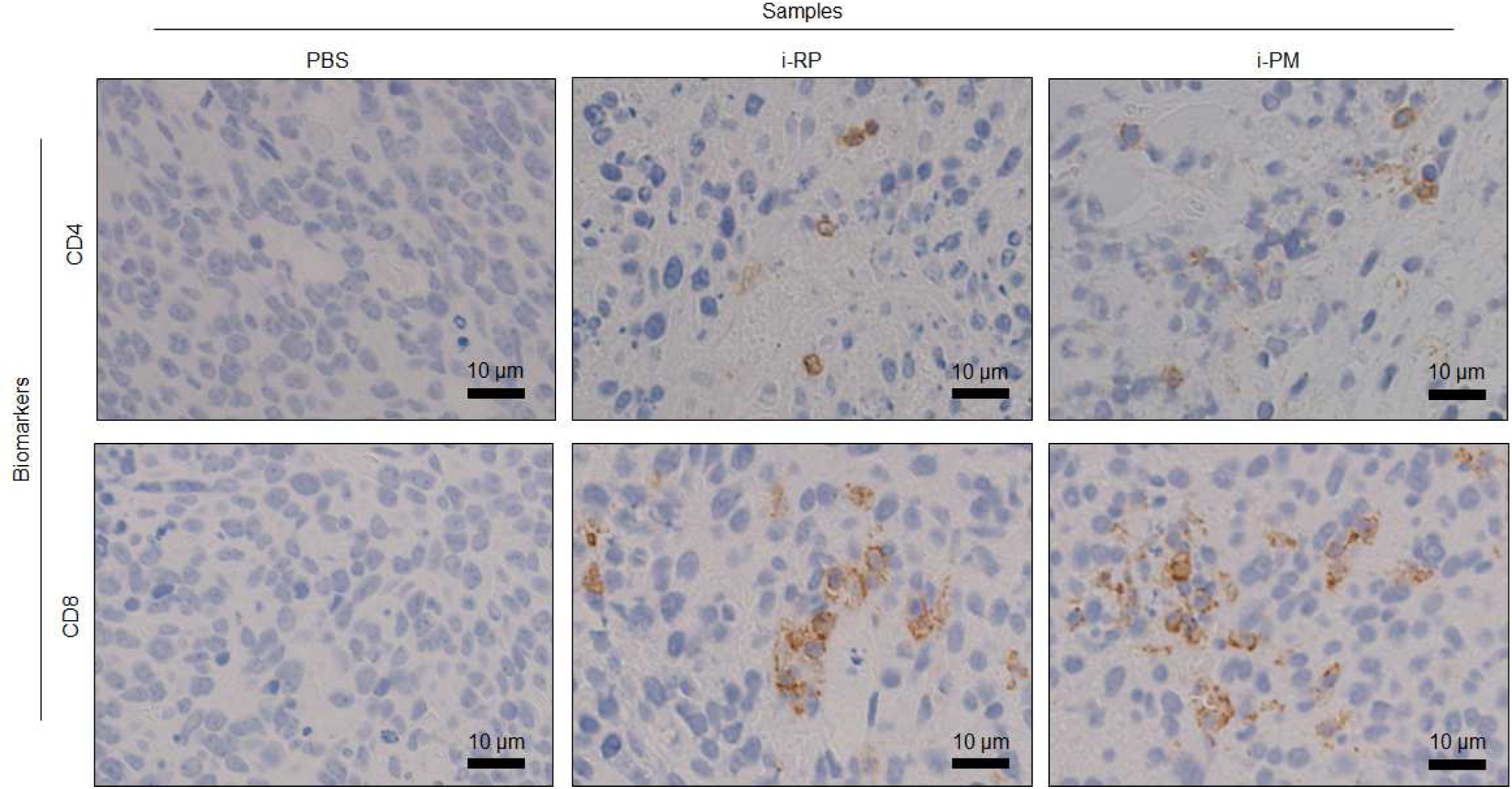
IHC (CD4 and CD8) stained tumour tissues collected from mice at day 1 after treatment with PBS, i-RP, and i-PM.

**Supplementary Figure S4.**
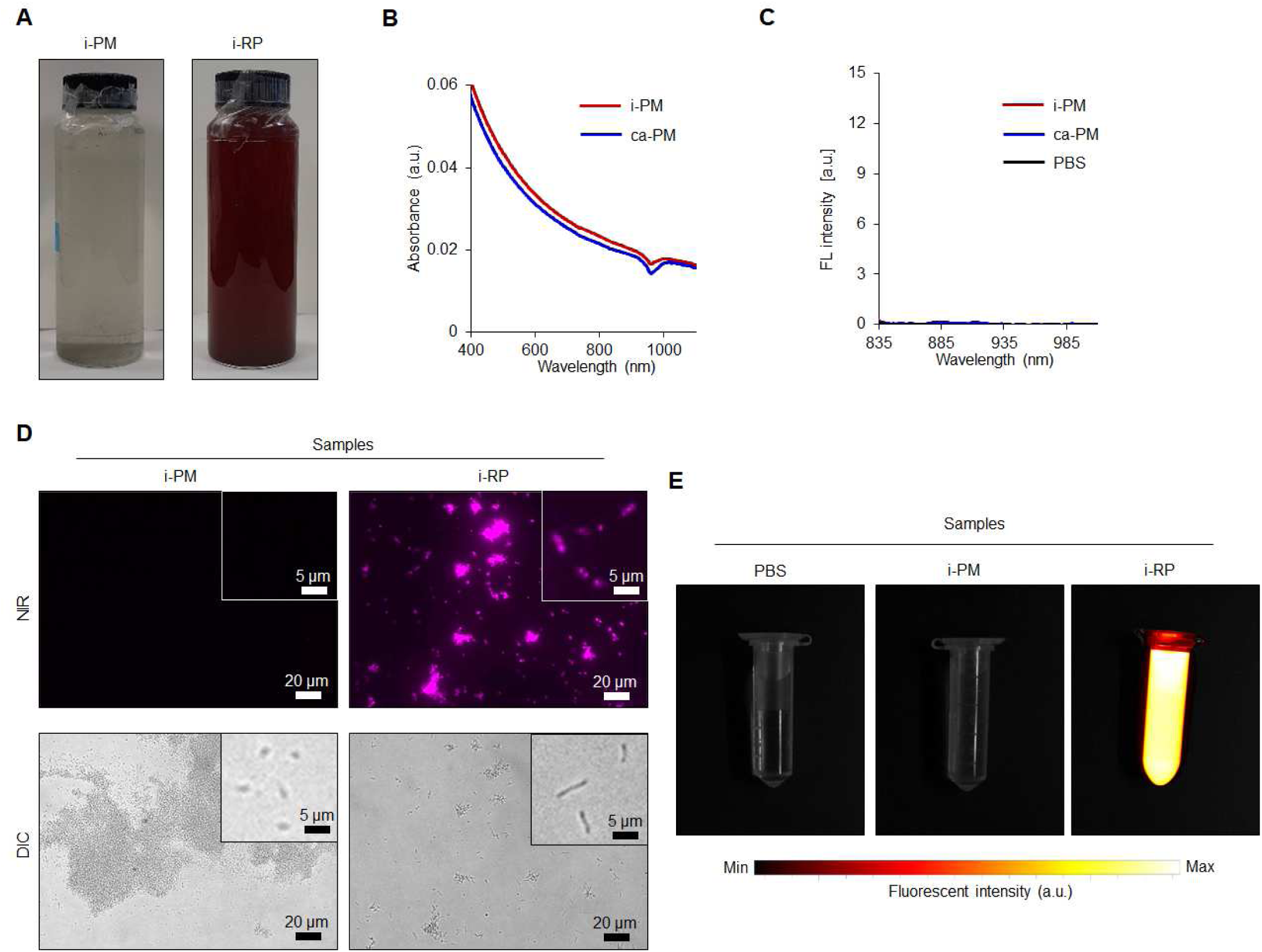
Optical properties of functional bacteria. (**A**) Images of i-PM (left) and i-RP (right) dispersions. (**B**) UV‒Vis‒NIR absorbance spectra of i-PM and ca-PM. (**C**) Fluorescent emission spectra of i-PM, ca-PM, and PBS excited at 805 nm. (**D**) *In vitro* NIR fluorescent and differential interference contrast (DIC) imaging of i-PM and i-RP. Bacterial concentration is 5 × 10^8^ CFU/mL. Upper-right inset of fluorescent/DIC images are the magnified i-PM and i-RP. The bacteria display a pink fluorescence. (**E**) NIR FL images of PBS, i-PM, and i-RP dispersions.

**Supplementary Figure S5.**
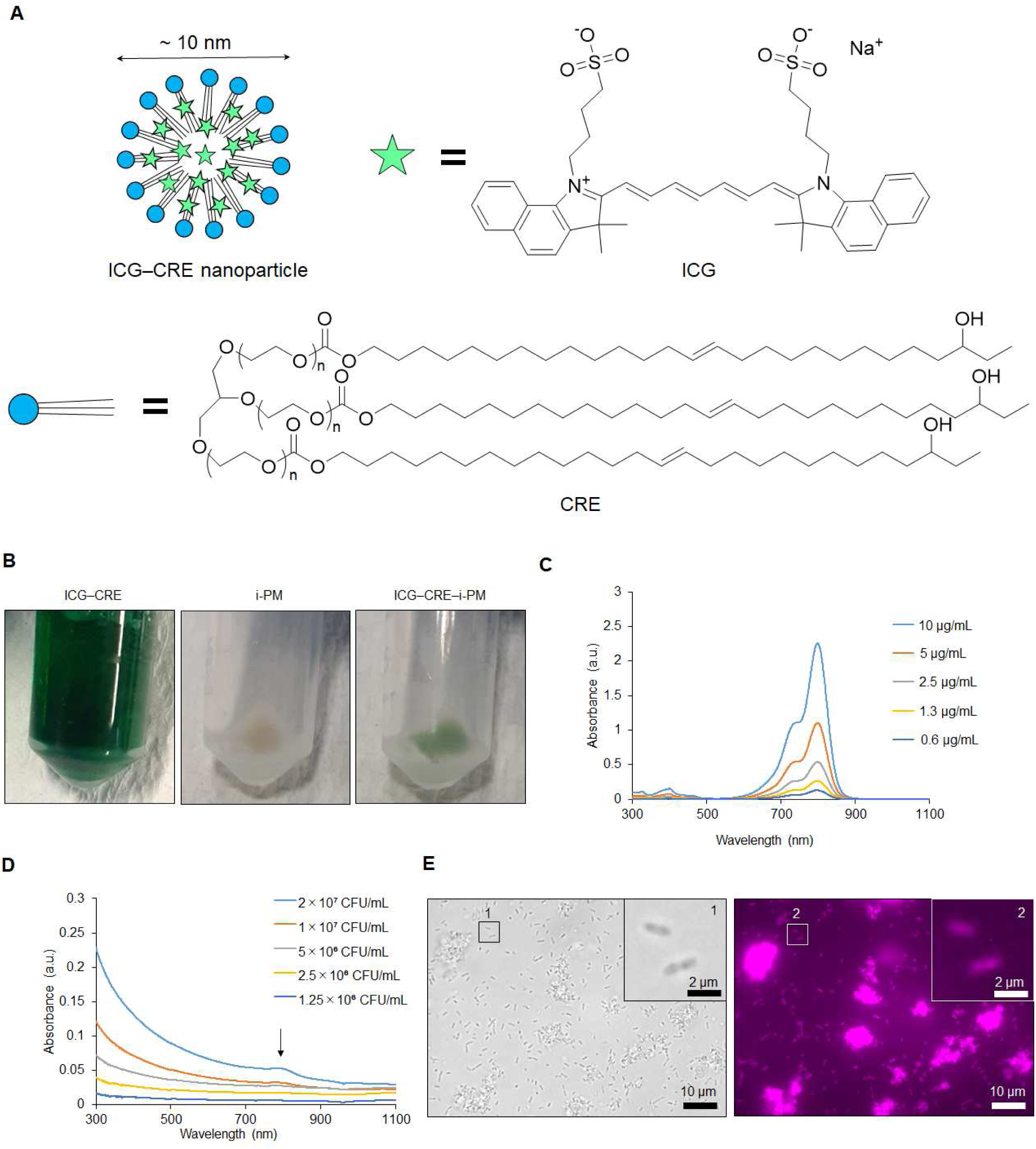
Optical properties of ICG‒CRE‒i-PM. (**A**) Schematic illustration of ICG‒CRE nanoparticle. (**B**) Images of ICG‒CRE dispersion (left), and i-PM (middle) and ICG‒CRE‒i-PM (right) pellets. (**C**) UV‒Vis‒NIR absorbance of ICG‒CRE dispersions at different concertation. (**D**) UV‒vis‒NIR absorbance spectra of ICG‒CRE‒BB dispersions at different concentrations. Black arrows display a characteristic peak derived from ICG molecule around at 800 nm. (**E**) DIC and NIR fluorescent imaging of ICG‒CRE‒i-PM prepared. The numbers (1 and 2) represent the location for magnified images of bacteria.

**Supplementary Figure S6.**
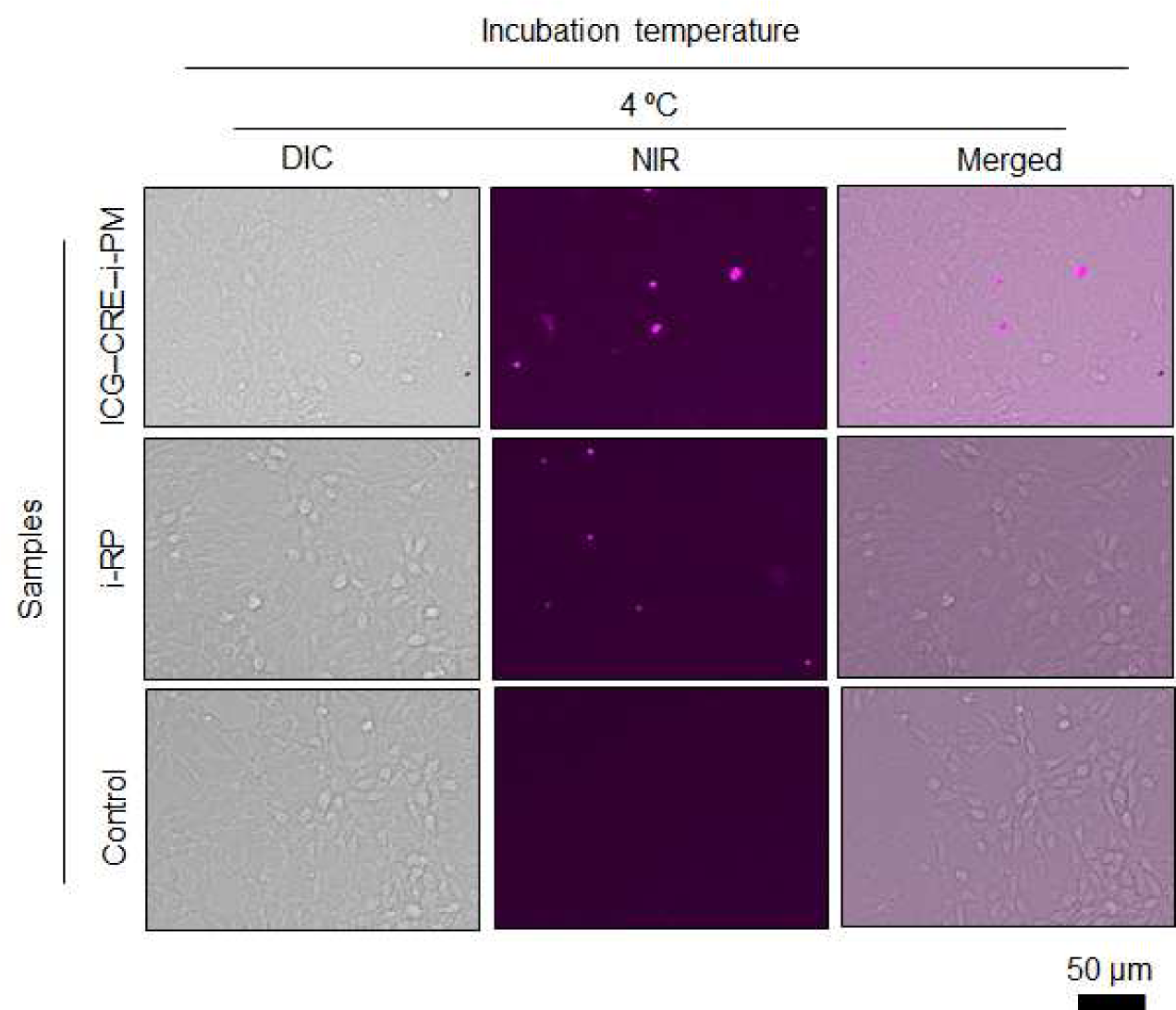
FL images of live Colon-26 cells after treatment with ICG‒CRE‒i-PM and i-RP for 4 h at 4 °C. The bacteria display a pink fluorescence.

**Supplementary Figure S7.**
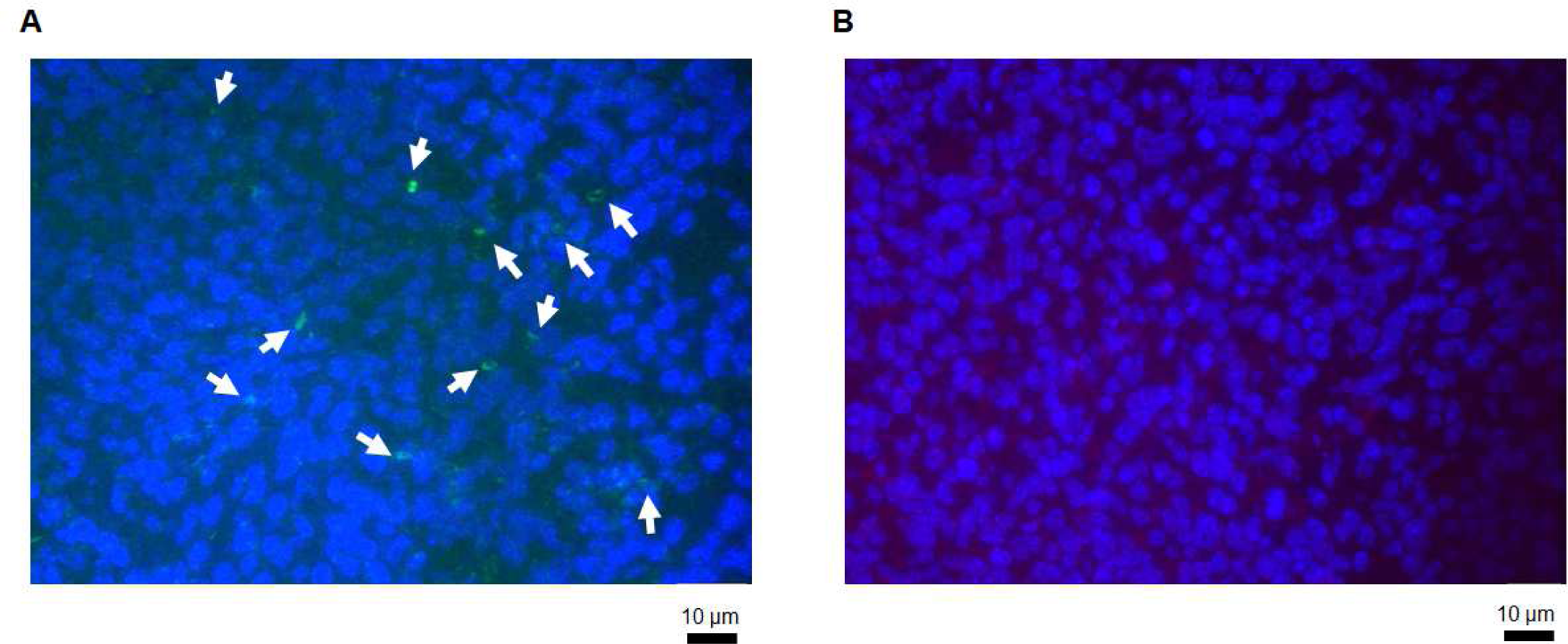
Observation of bacterial distribution in a solid tumour tissue using FISH analysis. Bacterial cells and colonies of (**A**) PM and (**B**) RP are coloured from green and red, respectively. Cancer cells (blue) was counterstained with 4’,6-diamidino-2-phenylindole (DAPI). White arrows represent tumour-resident PM colonies.

**Supplementary Figure S8.**
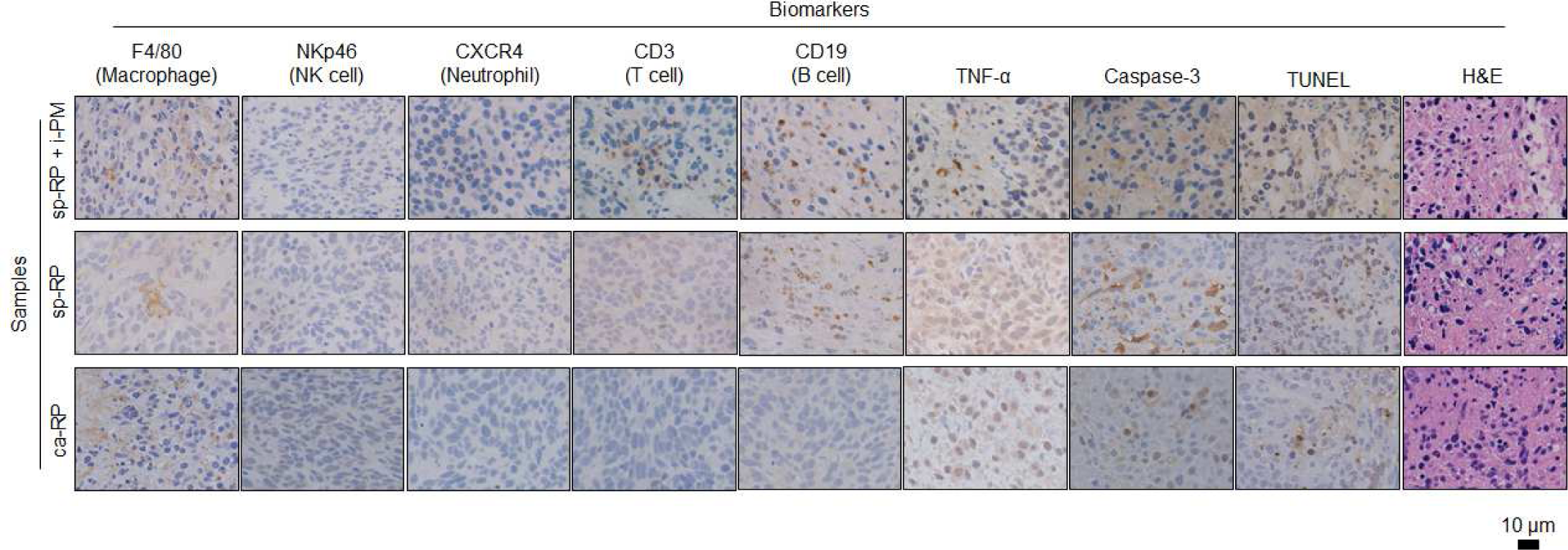
IHC (F4/80, NKp46, CXCR4, CD3, CD19, TNF-α and caspase-3), TUNEL, and H&E stained tumour tissues collected from the groups of mice at day 1 after treatments with ca-RP, sp-RP, and sp-RP + i-PM.

**Supplementary Figure S9.**
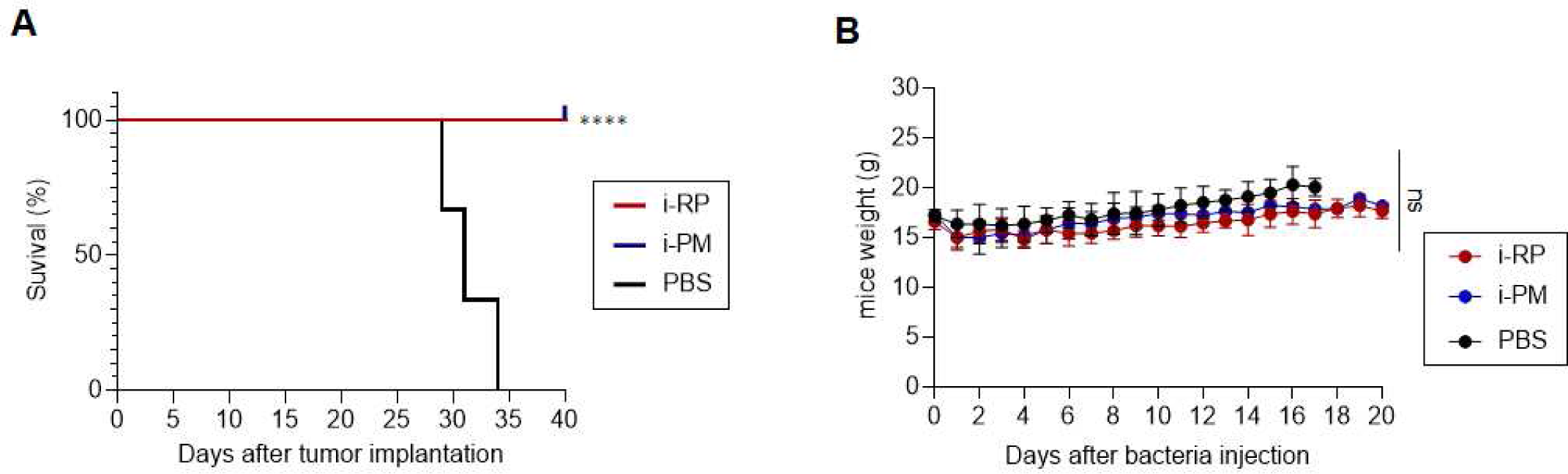
(**A**) Kaplan–Meier survival curves of sarcoma-bearing mice (n = 3 biologically independent mice) after tumour implantation for 40 days. Statistical significance was calculated in comparison with the PBS group. ****, *p* < 0.0001. (**B**) Weight of mice after each treatment. Data are represented as means ± SEM; n = 3 independent experiments. ns, not significant.

**Supplementary Figure S10.**
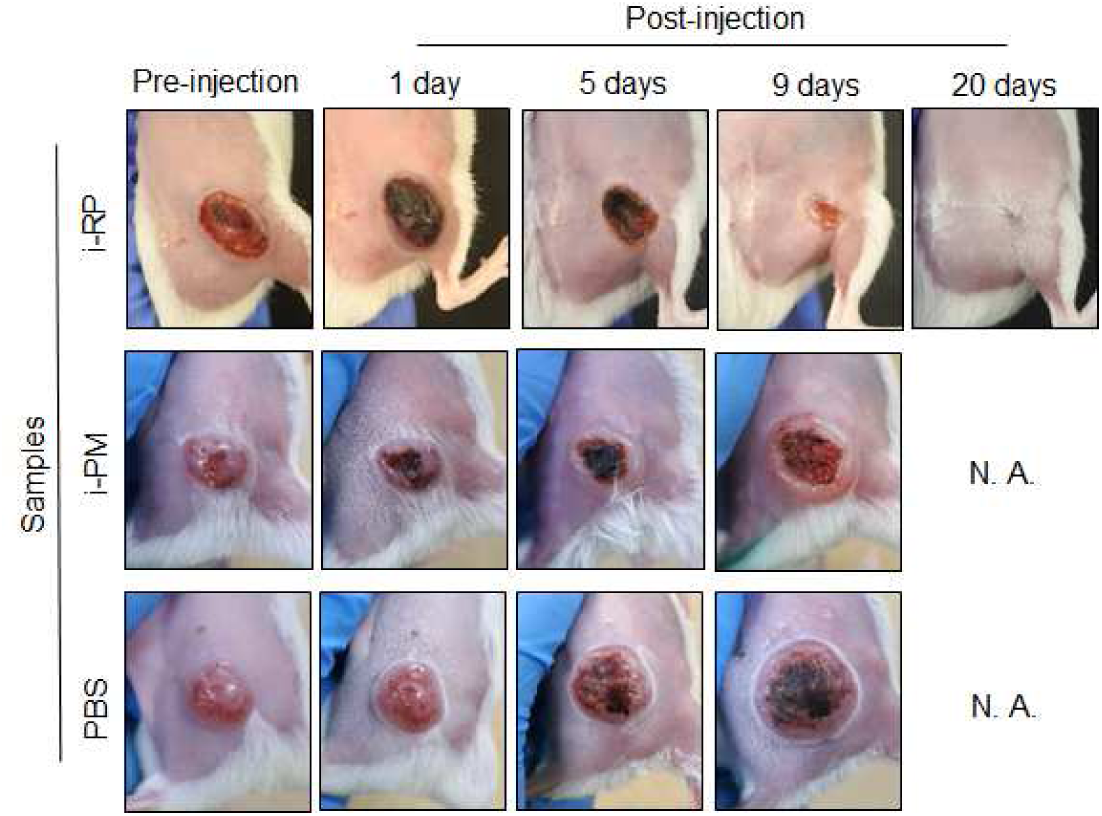
*In vivo* antitumor efficacy of functional bacteria against drug-resistant cancer model. Images of lungs after each treatment.

**Supplementary Figure S11.**
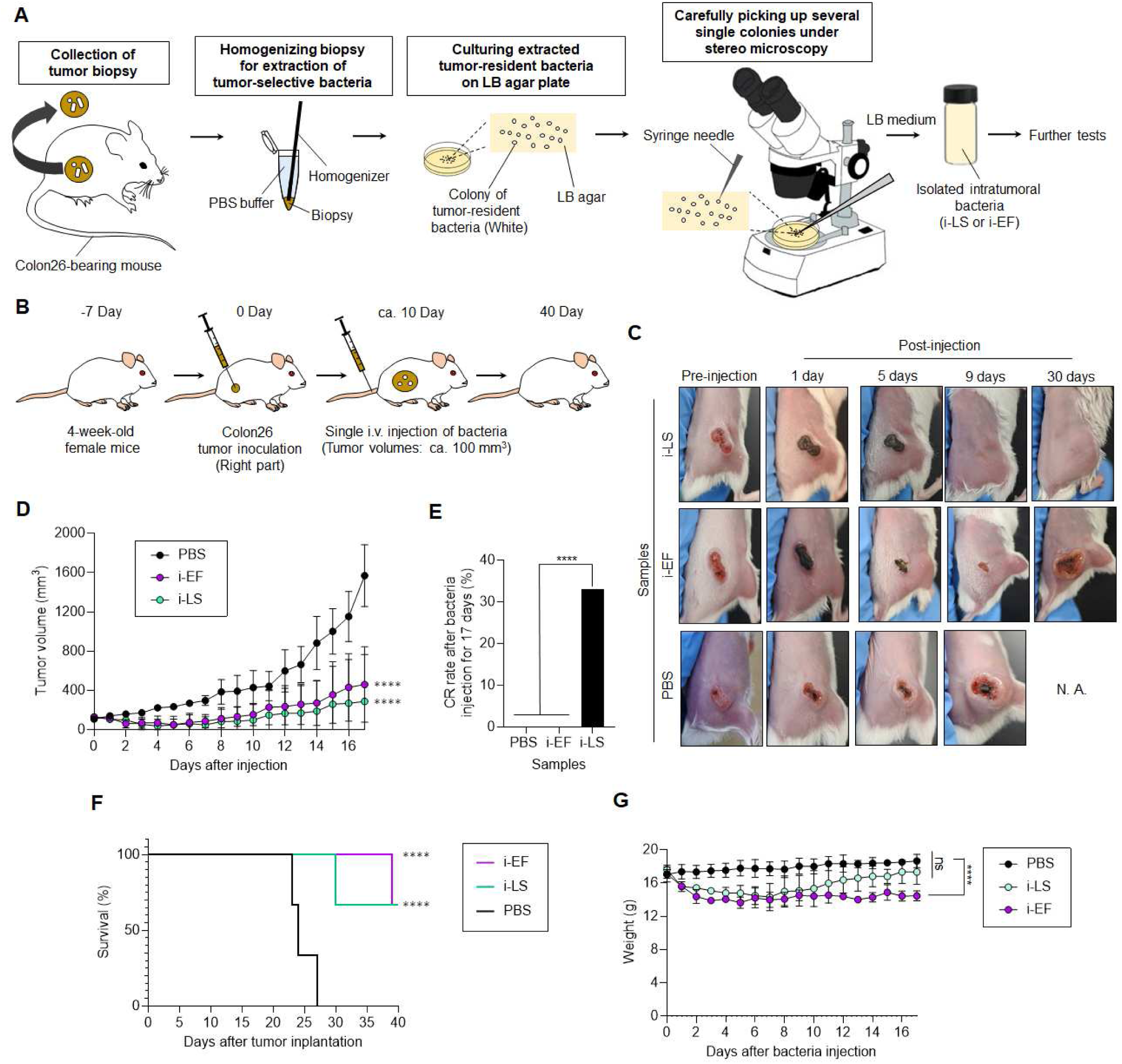
*In vivo* antitumour efficacy of various intramural bacteria. (**A**) Schematic illustration of isolations of i-LS and i-EF from solid tumours. (**B**) Schematic illustration of *in vivo* Colon-26 carcinoma antitumour tests using i-LS and i-EF. (**C**) Images of mice after each treatment. (**D**) *In vivo* anticancer effect of i-LS and i-EF. The PBS or bacterial suspension was intravenously injected into Colon26-bearing mice. Data are represented as mean ± SEM; n = 3 biologically independent mice. Statistical significance was calculated in comparison with the PBS group. ****, *p* < 0.0001. (**E**) CR rate of Colon-26-bearing mice (n = 3 biologically independent mice) after bacteria or PBS injection for 17 days with and without laser irradiation. ****, *p* < 0.0001. (**F**) Kaplan–Meier survival curves of Colon26-bearing mice (n = 3 biologically independent mice) after tumour implantation for 40 days. Statistical significance was calculated in comparison with PBS group. ****, *p* < 0.0001. (**G**) Weight of mice after each treatment. Data are represented as mean ± SEM; n = 3 independent experiments. ns, not significant. ****, *p* < 0.0001.

**Supplementary Table S1.**
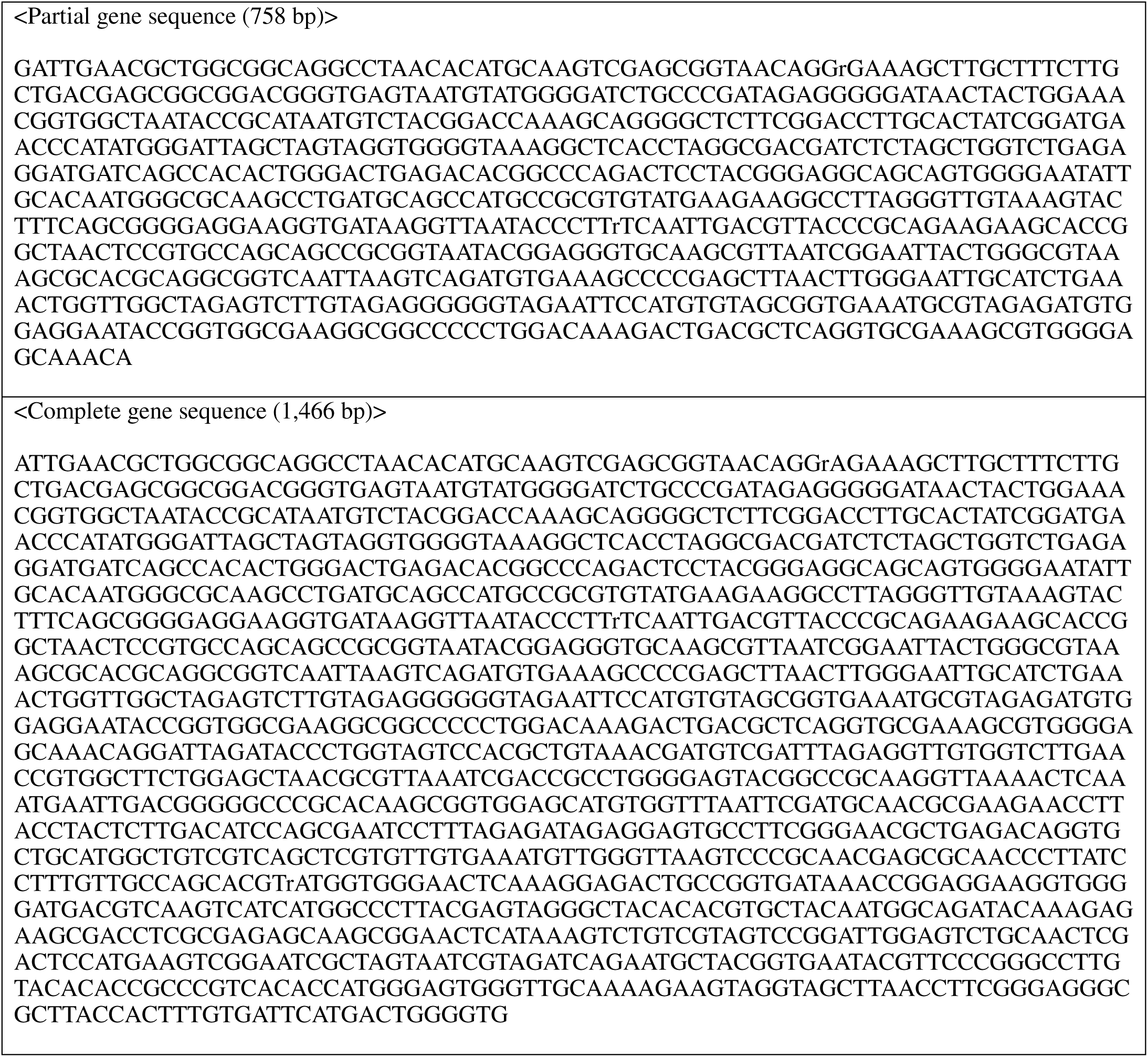
Obtained 16S rRNA gene sequences of i-PM.

**Supplementary Table S2.**
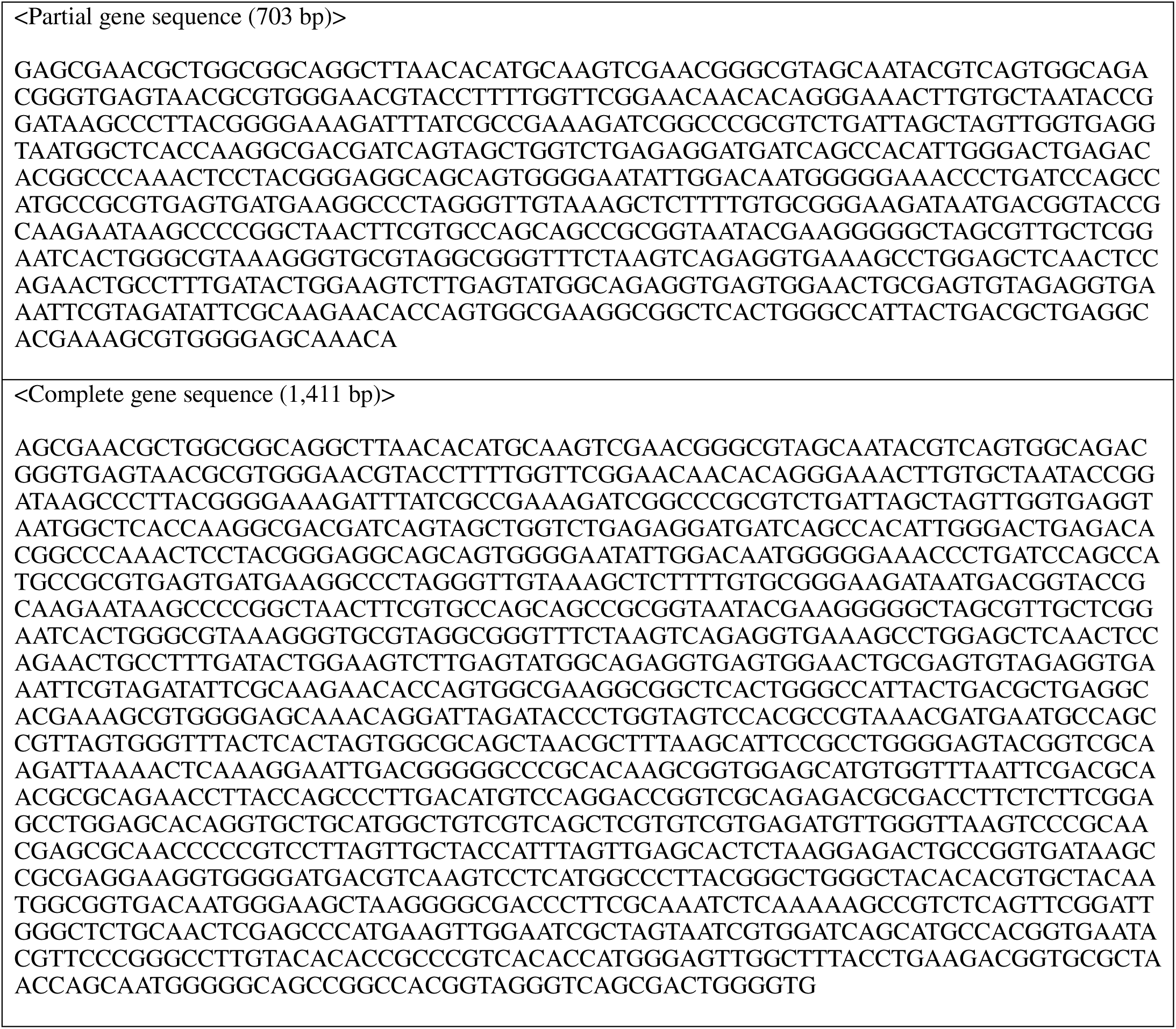
Obtained 16S rRNA gene sequences of sp-RP.

**Supplementary Table S3.** CBCs and biochemical parameters of the mice injected with PBS or i-RP dispersion after 30 days.

| Measured value | Entry | Unit | PBS (n = 5) | i-RP (n = 5) | P value |
| --- | --- | --- | --- | --- | --- |
| CBC | WBC | $\times 10^2 / \mu\text{L}$ | $56.00 \pm 7.07$ | $56.33 \pm 7.57$ | $> 0.05$ |
| | RBC | $\times 10^4 / \mu\text{L}$ | $865.80 \pm 19.25$ | $876.20 \pm 50.79$ | $> 0.05$ |
| | HGB | g/dL | $12.58 \pm 0.37$ | $12.66 \pm 0.81$ | $> 0.05$ |
| | HCT | % | $40.74 \pm 0.79$ | $40.82 \pm 2.40$ | $> 0.05$ |
| | MCV | fL | $47.08 \pm 0.71$ | $46.58 \pm 0.53$ | $> 0.05$ |
| | MCH | pg | $14.52 \pm 0.26$ | $14.44 \pm 0.15$ | $> 0.05$ |
| | MCHC | g/dL | $30.09 \pm 0.49$ | $31.00 \pm 0.33$ | $> 0.05$ |
| | PLT | $\times 10^4 / \mu\text{L}$ | $67.46 \pm 4.97$ | $68.10 \pm 5.05$ | $> 0.05$ |
| Biochemical parameters | TP | g/dL | $4.10 \pm 0.13$ | $4.00 \pm 0.18$ | $> 0.05$ |
| | ALB | g/dL | $2.90 \pm 0.09$ | $2.70 \pm 0.13$ | $> 0.05$ |
| | BUN | mg/dL | $21.02 \pm 3.61$ | $21.94 \pm 2.77$ | $> 0.05$ |
| | CRE | mg/dL | $0.10 \pm 0.01$ | $0.10 \pm 0.01$ | $> 0.05$ |
| | Na | mEq/L | $151.40 \pm 1.34$ | $152.40 \pm 1.52$ | $> 0.05$ |
| | K | mEq/L | $3.38 \pm 0.28$ | $3.14 \pm 0.36$ | $> 0.05$ |
| | Cl | mEq/L | $118.20 \pm 1.10$ | $118.80 \pm 0.84$ | $> 0.05$ |
| | AST | IU/L | $64.00 \pm 8.49$ | $67.60 \pm 11.37$ | $> 0.05$ |
| | ALT | IU/L | $34.50 \pm 4.95$ | $35.40 \pm 3.98$ | $> 0.05$ |
| | LDH | IU/L | $284.25 \pm 60.32$ | $264.80 \pm 56.14$ | $> 0.05$ |
| | AMY | IU/L | $1806.00 \pm 157.14$ | $1853.20 \pm 193.67$ | $> 0.05$ |
| | CK | IU/L | $211.50 \pm 50.35$ | $205.67 \pm 71.46$ | $> 0.05$ |

Data are represented as the mean ± SEM; n = 5 biologically independent mice. Statistical analyses were performed using the Student's two-sided t-test.
Abbreviations: ALB, albumin; ALT, alanine transaminase; AMY, amylase; AST, aspartate aminotransferase; BUN, blood urea nitrogen; Cl, chlorine; CK, creatine kinase; CRE, creatinine; CRP, C-reactive protein; HCT, hematocrit; HGB, hemoglobin; K, potassium; LDH, lactate dehydrogenase; MCH, mean corpuscular hemoglobin; MCHC, mean corpuscular hemoglobin concentration; MCV, mean corpuscular volume; Na, sodium; PLT, platelet; RBC, red blood cell; TP, total protein; WBC, white blood cell.

**Supplementary Table S4.** CBCs and biochemical parameters of the mice injected with PBS or i-PM dispersion after 30 days.

| Measured value | Entry | Unit | PBS (n = 5) | i-PM (n = 5) | P value |
| --- | --- | --- | --- | --- | --- |
| CBC | WBC | $\times 10^2 / \mu\text{L}$ | $66.2 \pm 12.70$ | $66.8 \pm 12.13$ | $> 0.05$ |
| | RBC | $\times 10^4 / \mu\text{L}$ | $948.8 \pm 8.70$ | $883.8 \pm 31.58$ | $> 0.05$ |
| | HGB | g/dL | $14.1 \pm 0.31$ | $13.88 \pm 0.46$ | $> 0.05$ |
| | HCT | % | $44.7 \pm 0.91$ | $40.98 \pm 1.67$ | $> 0.05$ |
| | MCV | fL | $47.1 \pm 0.81$ | $46.38 \pm 0.27$ | $> 0.05$ |
| | MCH | pg | $14.8 \pm 0.19$ | $15.7 \pm 0.12$ | $> 0.05$ |
| | MCHC | g/dL | $31.5 \pm 0.40$ | $33.84 \pm 0.24$ | $> 0.05$ |
| | PLT | $\times 10^4 / \mu\text{L}$ | $72.6 \pm 4.67$ | $70.76 \pm 6.38$ | $> 0.05$ |
| Biochemical parameters | TP | g/dL | $4.1 \pm 0.15$ | $3.9 \pm 0.16$ | $> 0.05$ |
| | ALB | g/dL | $2.7 \pm 0.15$ | $2.6 \pm 0.06$ | $> 0.05$ |
| | BUN | mg/dL | $21.5 \pm 2.04$ | $22.0 \pm 1.85$ | $> 0.05$ |
| | CRE | mg/dL | $0.13 \pm 0.01$ | $0.12 \pm 0.03$ | $> 0.05$ |
| | Na | mEq/L | $153.0 \pm 1.14$ | $151.0 \pm 0.55$ | $> 0.05$ |
| | K | mEq/L | $3.6 \pm 0.42$ | $3.6 \pm 0.14$ | $> 0.05$ |
| | Cl | mEq/L | $117.2 \pm 0.45$ | $117.8 \pm 1.30$ | $> 0.05$ |
| | AST | IU/L | $51.4 \pm 6.43$ | $58.8 \pm 10.43$ | $> 0.05$ |
| | ALT | IU/L | $33.2 \pm 7.36$ | $34.6 \pm 6.58$ | $> 0.05$ |
| | LDH | IU/L | $244.2 \pm 75.88$ | $254.6 \pm 68.68$ | $> 0.05$ |
| | AMY | IU/L | $1671.4 \pm 97.66$ | $1682.8 \pm 131.58$ | $> 0.05$ |
| | CK | IU/L | $181.6 \pm 113.95$ | $182.6 \pm 128.41$ | $> 0.05$ |

Data are represented as the mean ± SEM; n = 5 biologically independent mice. Statistical analyses were performed using the Student's two-sided t-test.
Abbreviations: ALB, albumin; ALT, alanine transaminase; AMY, amylase; AST, aspartate aminotransferase; BUN, blood urea nitrogen; Cl, chlorine; CK, creatine kinase; CRE, creatinine; CRP, C-reactive protein; HCT, hematocrit; HGB, hemoglobin; K, potassium; LDH, lactate dehydrogenase; MCH, mean corpuscular hemoglobin; MCHC, mean corpuscular hemoglobin concentration; MCV, mean corpuscular volume; Na, sodium; PLT, platelet; RBC, red blood cell; TP, total protein; WBC, white blood cell.

**Supplementary Table S5.**
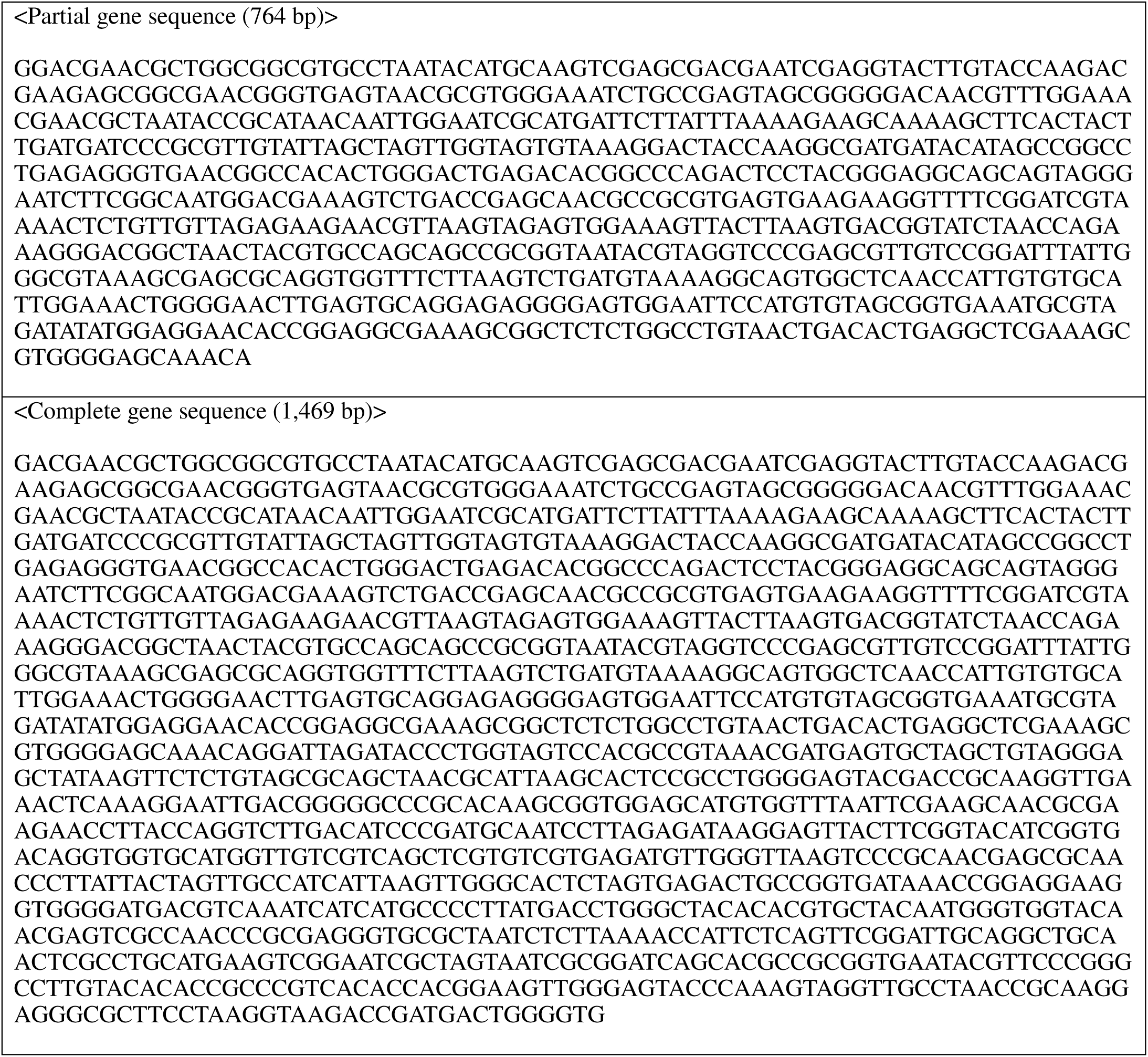
Obtained 16S rRNA gene sequences of i-LS.

**Supplementary Table S6.**
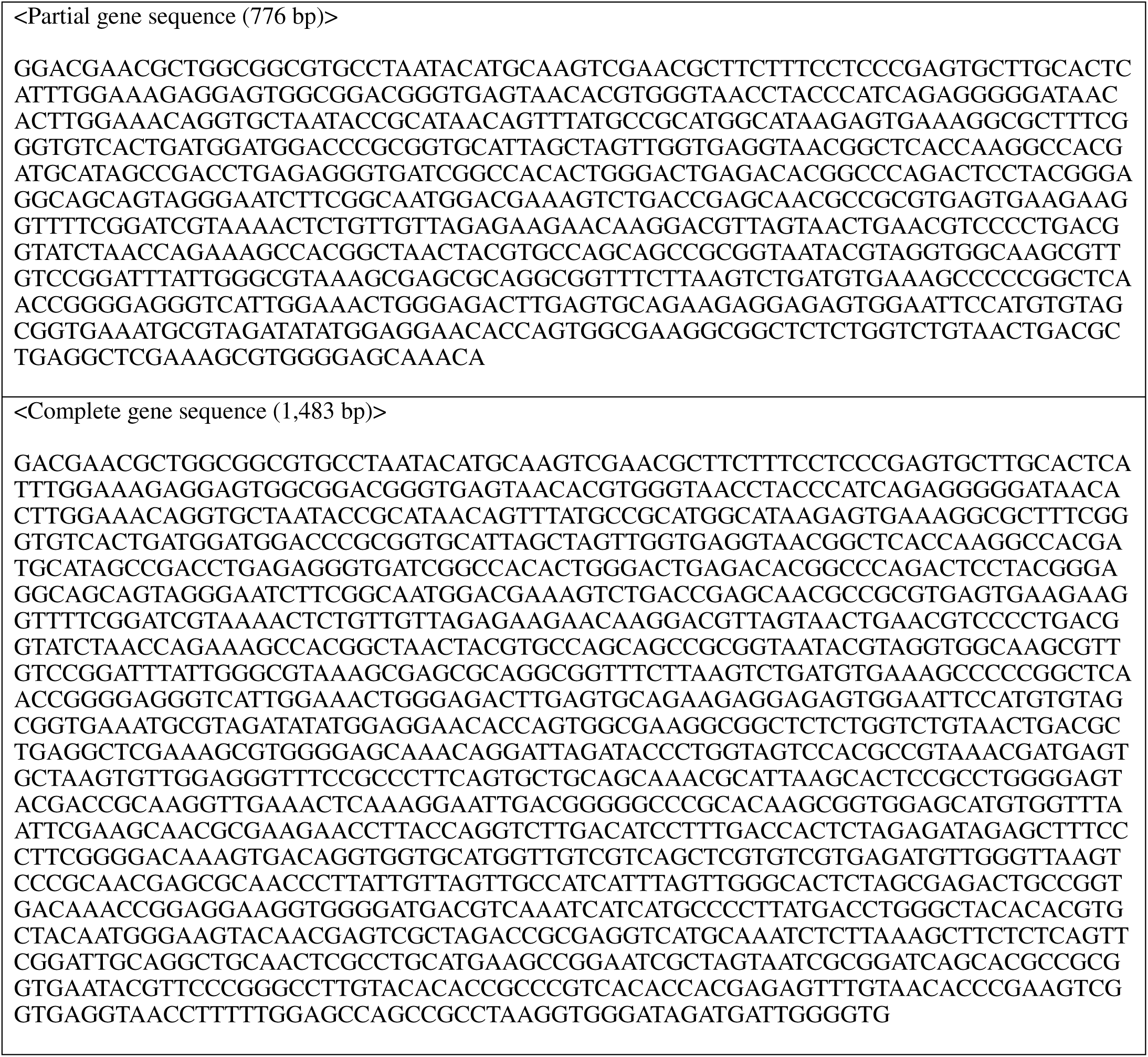
Obtained 16S rRNA gene sequences of i-EF

**Supplementary Table S7.**
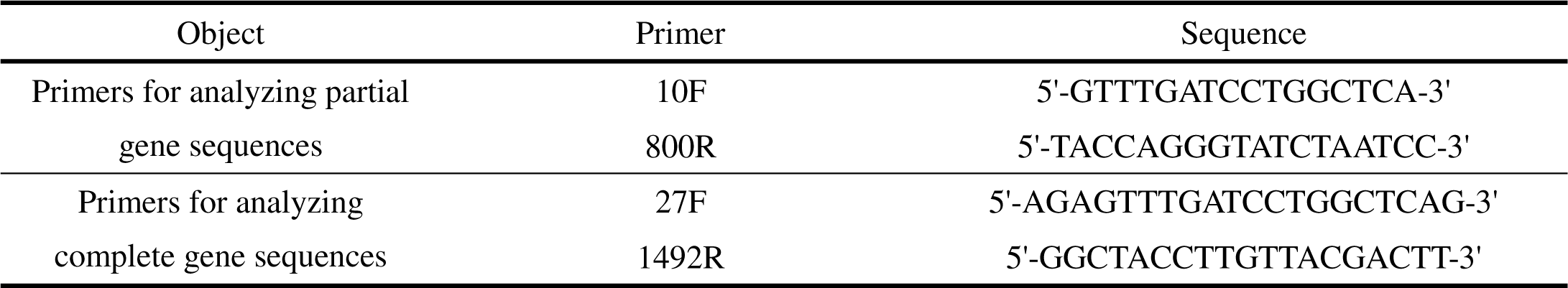
Primers used for amplification and sequencing of the 16S rRNA gene of i-PM and i-RP.

**Supplementary Table S8.**
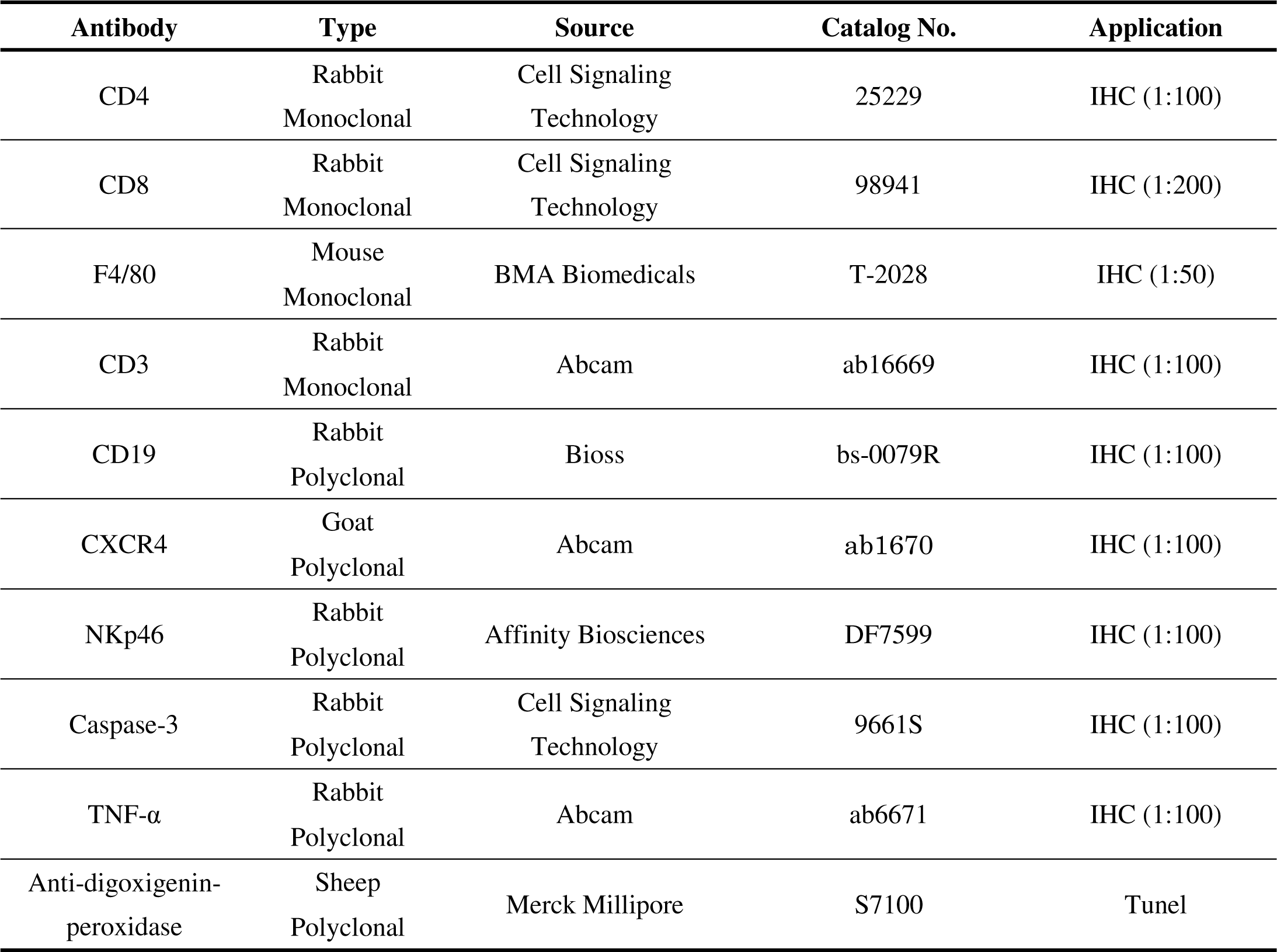
Antibodies used in this study.

**Supplementary Table S9.**
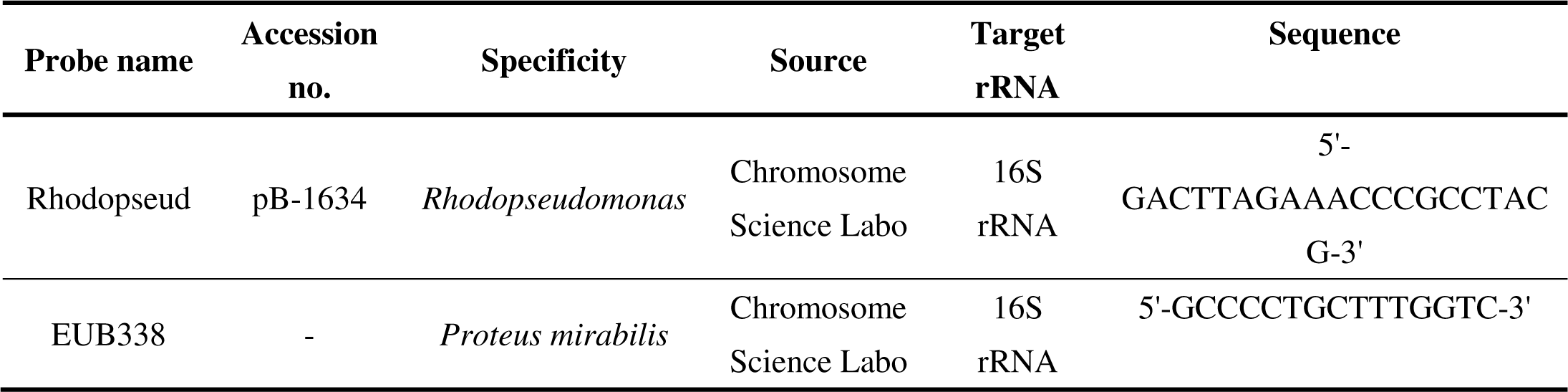
Probes used in microbial FISH analysis.

## References

1. Kumar, N., Fazal, S., Miyako, E., Matsumura, K. & Rajan, R. Avengers against cancer: A new era of nano-biomaterial-based therapeutics. Materials Today 51, 317–349 (2021).

2. Bacterial therapy of cancer: Methods and protocols (ed Hoffman, R. M.) (Humana Press, Springer, Netherlands) (2015).

3. Zhou, S., Gravekamp, C., Bermudes, D. & Liu, K. Tumor-targeting bacteria engineered to fight cancer. Nat. Rev. Cancer 18, 727–743 (2018).

4. Forbes, N. S. Engineering the perfect (bacterial) cancer therapy. Nat. Rev. Cancer 10, 785–794 (2010).

5. Brown, J. M. & William, W. R. Exploiting tumor hypoxia in cancer treatment. Nat. Rev. Cancer 4, 437–447 (2004).

6. Duong, M. T-Q., Qin, Y., You, S-H. & Min, J-J. Bacteria-cancer interactions: bacteria-based cancer therapy. Exp. Mol. Med. 51, 1–15 (2019).

7. Din, M., Danino, T., Prindle, A., Skalak, M., Selimkhanov, J., Allen, K., Julio, E., Atolia, E., Tsimring, L. S., Bhatia, S. N. & Hasty, J. Synchronized cycles of bacterial lysis for *in vivo* delivery. Nature 536, 81–85 (2016).

8. Bourdeau, R. W., Lee-Gosselin, A., Lakshmanan, A., Farhadi, A., Kumar, S. R., Nety, S. P. & Shapiro M. G. Acoustic reporter genes for noninvasive imaging of microorganisms in mammalian hosts. Nature 553, 86–90 (2018).

9. Chowdhury, S., Castro, S., Coker, C., Hinchliffe, T. E., Arpaia, N. & Danino, T. Programmable bacteria induce durable tumor regression and systemic antitumor immunity. Nat. Med. 25, 1057– 1063 (2019).

10. Harimoto, T., Hahn, J., Chen, Y., Im, J., Zhang, J., Hou, N., Coker, C., Gray, K., Harr, N., Chowdhury, C., Pu, K., Nimura, C., Arpaia, N., Leong, K. & Danino, T. A programmable encapsulation system improves delivery of therapeutic bacteria in mice. Nat. Biotechnol. (2022). https://doi.org/10.1038/s41587-022-01244-y

11. Felfoul, O., Mohammadi, M., Taherkhani, S., de Lanauze, D., Xu, Y. Z., Loghin, D., Essa, S., Jancik, S., Houle, D., Lafleur, M., Gaboury, L., Tabrizian, M., Kaou, N., Atkin, M., Vuong, T., Batist, G., Beauchemin, N., Radzioch, D. & Martel, S. Magneto-aerotactic bacteria deliver drug-containing nanoliposomes to tumor hypoxic regions. Nat. Nanotechnol. 11, 941–947 (2016).

12. Zheng, D., Chen, Y., Li, Z., Xu, L., Li, C., Li, B., Fan, J., Cheng, S. & Zhang, X. Optically-controlled bacterial metabolite for cancer therapy. Nat. Commun. 9, 1680 (2018).

13. Suh, S., Jo, A., Traore, M. A., Zhan, Y., Coutermarsh-Ott, S. L., Ringel-Scaia, V. M., Allen, I. C., Davis, R. M., Behkam, B. Nanoscale bacteria-enabled autonomous drug delivery system (NanoBEADS) enhances intratumoral transport of nanomedicine. Adv. Sci. 6, 1801309 (2019).

14. Yang, X., Komatsu, S., Reghu, S. & Miyako, E. Optically activatable photosynthetic bacteria-based highly tumor specific immunotheranostics. Nano Today 37, 101100 (2021).

15. Kalaora, S., Nagler, A., Nejman, D., Alon, M., Barbolin, C., Barnea, E., Ketelaars, S. L. C., Cheng, K., Vervier, K., Shental, N., Bussi, Y., Rotkopf, R., Levy, R., Benedek, G., Trabish, S., Dadosh, T., Levin-Zaidman, S., Geller, L. T., Wang, K., Greenberg, P., Yagel, G., Peri, A., Fuks, G., Bhardwaj, N., Reuben, A., Hermida, L., Johnson, S. B., Galloway-Peña, J. R., Shropshire, W. C., Bernatchez, C., Haymaker, C., Arora, R., Roitman, L., Eilam, R., Weinberger, A., Lotan-Pompan, M., Lotem, M., Admon, A., Levin, Y., Lawley, T. D., Adams, D. J., Levesque, M. P., Besser, M. J., Schachter, J., Golani, O., Segal, E., Geva-Zatorsky, N., Ruppin, R., Kvistborg, P., Peterson, S. N., Wargo, J. A., Straussman, R., Samuels, Y. Identification of bacteria-derived HLA-bound peptides in melanoma. Nature 592, 138–143 (2021).

16. Sepich-Poore, G. D., Zitvogel, L., Straussman, R., Hasty, J., Wargo, J. A., Knight, R. The microbiome and human cancer. Science 371, eabc4552 (2021).

17. Nejman, D., Livyatan, I., Fuks, G., Gavert, N., Zwang, Y., Geller, L. T., Rotter-Maskowitz, A., Weiser, R., Mallel, G., Gigi, E., Meltser, A., Douglas, G. M., Kamer, I., Gopalakrishnan, V., Dadosh, T., Levin-Zaidman, S., Avnet, S., Atlan, T., Cooper, Z. A., Arora, R., Cogdill, A. P., Khan, M. A. W., Ologun, G., Bussi, Y., Weinberger, A., Lotan-Pompan, M., Golani, O., Perry, G., Rokah, M., Bahar-Shany, K., Rozeman, E. A., Blank, C. U., Ronai, A., Shaoul, R., Amit, A., Dorfman, T., Kremer, R., Cohen, Z. R., Harnof, S., Siegal, T., Yehuda-Shnaidman, E., Gal-Yam, E. N., Shapira, H., Baldini, N., Langille, M. G. I., Ben-Nun, A., Kaufman, B., Nissan, A., Golan, T., Dadiani, M., Levanon, K., Bar, J., Yust-Katz, S., Barshack, I., Peeper, D. S., Raz, D. J., Segal, E., Wargo, J. A., Sandbank, J., Shental, N. & Straussman, R. The human tumor microbiome is composed of tumor type-specific intracellular bacteria. Science 368, 973–980 (2020).

18. Geller, L. T., Barzily-Rokni, M., Danino, T., Jonas, O. H., Shental, N., Nejman, D., Gavert, N., Zwang, Y., Cooper, Z. A., Shee, K., Thaiss, C. A., Reuben, A., Livny, J., Avraham, R., Frederick, D. T., Ligorio, M., Chatman, K., Johnston, S. E., Mosher, C. M., Brandis, A., Fuks, G., Gurbatri, C., Gopalakrishnan, V., Kim, M., Hurd, M. W., Katz, M., Fleming, J., Maitra, A., Smith, D. A., Skalak, M., Bu, J., Michaud, M., Trauger, S. A., Barshack, I., Golan, T., Sandbank, J., Flaherty, K. T., Mandinova, A., Garrett, W. S., Thayer, S. P., Ferrone, C. R., Huttenhower, C., Bhatia, S. N., Gevers, D., Wargo, J. A., Golub, T. R. & Straussman, R. Potential role of intratumor bacteria in mediating tumor resistance to the chemotherapeutic drug gemcitabine. Science 357, 1156–1160 (2017).

19. Fu, A., Yao, B., Dong, T., Chen, Y., Yao, J., Liu, Y. Li, H., Bai, H., Liu, X., Zhang, Y., Wang, C., Guo, Y., Li, N. & Cai, S. Tumor-resident intracellular microbiota promotes metastatic colonization in breast cancer. Cell 185, 1356–1372 (2022).

20. Proteus mirabilis: Methods and protocols (ed Pearson, M. M.) (Humana Press, Springer, Netherlands) (2015).

21. Liu, C-J. Zhang, Y-L., Shang, Y., Wu, B., Yang, E., Luo, Y-Y., Li, X-R. Intestinal bacteria detected in cancer and adjacent tissue from patients with colorectal cancer. Oncol. Lett. 17, 1115–1127 (2015).

22. D’Mello, A. & Yotis, W. W. The action of sodium deoxycholate on *Escherichia coli*. Appl. Environ. Microbiol. 53, 1944–1946 (1987).

23. McCarty, N. S. & Ledesma-Amaro, R. Synthetic biology tools to engineer microbial communities for biotechnology. Trends Biotechnol. 37, 181–197 (2019).

24. Grosskopf, T. & Soyer, O. S. Synthetic microbial communities. Curr. Opin. Microbiol. 18, 72–77 (2014).

25. Zhang, H., Diao, H., Jia, L., Yuan, Y., Thamm, D. H., Wang, H., Jin, Y., Pei, S., Zhou, B., Yu, F., Zhao, L., Cheng, N., Du, H., Huang, Y., Zhang, D. & Lin, D. *Proteus mirabilis* inhibits cancer growth and pulmonary metastasis in a mouse breast cancer model. PLoS One 12, e0188960 (2017).

26. Netea, M. G., Domínguez-Andrés, J., Barreiro, L. B., Chavakis, T., Divangahi, M., Fuchs, E., Joosten, L. A. B., van der Meer, J. W. N., Mhlanga, M. M., Mulder, W. J. M., Riksen, N. P., Schlitzer, A., Schultze, J. L., Benn, C. S., Sun, J. C., Xavier, R. J. & Latz, E. Defining trained immunity and its role in health and disease. Nat. Rev. Immunol. 20, 375–388 (2020).

27. Mueller, S. N. & Mackay, L. K. Tissue-resident memory T cells: local specialists in immune defence. Nat. Rev. Immunol. 16, 79–89 (2016).

28. The Purple Phototrophic Bacteria (eds Hunter CN, Daldal F, Thurnauer MC, Beatty JT.) (Springer, Netherlands) (2009).

29. Zhao, M., Yang, M., Li, X-M., Jiang, P., Baranov, E., Li, S., Xu, M., Penman, S. & Hoffman, R.M. Tumor-targeting bacterial therapy with amino acid auxotrophs of GFP-expressing *Salmonella typhimurium*. Proc. Natl. Acad. Sci. USA 102, 755–760 (2005).

30. Zhao, M., Yang, M., Ma, H., Li, X., Tan, X., Li, S., Yang, Z. & Hoffman, R. M. Targeted therapy with a *Salmonella typhimurium* leucine-arginine auxotroph cures orthotopic human breast tumors in nude mice. Cancer Res. 66, 7647–7652 (2006).

31. Reghu, S. & Miyako, E. Nanoengineered *Bifidobacterium bifidum* with optical activity for photothermal cancer immunotheranostics. Nano Lett. 22, 1880–1888 (2022).

32. Cossart, P. & Helenius, A. Endocytosis of viruses and bacteria. Cold Spring Harb. Perspect. Biol. 6, a016972 (2014).

33. Fluorescence in situ hybridization (FISH) (ed Liehr, T.) (Springer, Netherlands) (2010).

34. Stackebrandt, E. & Ebers, J. Taxonomic parameters revisited: tarnished gold standards. Microbiol. Today 33, 152–155 (2006).

35. Prasad, V., De Jesús, K. & Mailankody, S. The high price of anticancer drugs: origins, implications, barriers, solutions. Nat. Rev. Clin. Oncol. 14, 381–390 (2017).

36. Impact of COVID-19 on people’s livelihoods, their health and our food systems (Joint statement by ILO, FAO, IFAD and WHO) (2020).

37. Synthetic biology: Parts, devices and applications (ed Smolke, C., et al.) (Wiley, New Jersey, US) (2018).

38. Zhang, J., Hoedt, E. C., Liu, Q., Berendsen, E., Teh, J. J., Hamilton, A., O’ Brien, A. W., Ching, J. Y. L., Wei, H., Yang, K., Xu, Z., Wong, S. H., Mak, J. W. Y., Sung, J. J. Y., Morrison, M., Yu, J., Kamm, M. A. & Ng, S. W. Elucidation of *Proteus mirabilis* as a key bacterium in crohn’s disease inflammation. Gastroenterology 160, 317–330 (2021).

